# How do sapling and adult demography explain beech predominance along environmental gradients in Central European forests?

**DOI:** 10.1101/2023.11.23.568436

**Authors:** Lukas Heiland, Georges Kunstler, Lisa Hülsmann

**Affiliations:** University of Bayreuth, Bayreuth Center of Ecology and Environmental Research (BayCEER), Ecosystem Analysis and Simulation (EASI) Lab, Dr.-Hans-Frisch-Str. 1–3, 95448 Bayreuth, Germany. Theoretical Ecology, Universität Regensburg, Universitätsstraße 31, 93053 Regensburg, Germany; Université Grenoble Alpes, Inrae, LESSEM, Grenoble, France; University of Bayreuth, Bayreuth Center of Ecology and Environmental Research (BayCEER), Ecosystem Analysis and Simulation (EASI) Lab, Germany

**Keywords:** dynamic species distribution model, Bayesian inverse calibration, ecogram, Fagus sylvatica, German national forest inventory, competition response, shade tolerance, Ellenberg theory

## Abstract

Understanding how species abundances are driven by biotic interactions along environmental gradients is a fundamental question in ecology. For abundances at competitive equilibria in Central European forests, a classical ecological theory formulated by Ellenberg (1963) predicts that beech (*Fagus sylvatica* L.) outcompetes other tree species within a mesic range of soil pH and water levels, while other species prevail under less favorable conditions. While the theory is generally accepted in forest ecology, only certain aspects of it have been substantiated by empirical evidence. Moreover, the demographic processes driving the turnover from beech to other tree species at the extremes of the soil gradients remain mostly unexplained.

To address this, we inversely calibrated a parsimonious forest model (JAB model) with a sapling stage and interacting populations with short time series of observed tree abundances from the German national forest inventory. By modelling how demographic rates vary along pH and soil water gradients, we were able to test the prediction that beech naturally predominates only at favorable soil conditions. Moreover, we tested with simulations how the environmental response of demographic rates explains *Fagus*’ changing relative abundance along the soil gradients.

Our results largely confirm that *Fagus* out competes other species in a central environmental range. Environmental change of *Fagus*’ relative abundance is primarily explained by environmental variation of its net basal area increment, followed by its competition response at the overstory and at the sapling stage. We found that even though sapling tolerance to shading is the primary mechanism for *Fagus* predominance, it only plays a secondary role for the environmental variation of its relative abundance.

*Synthesis:* By inverse calibration of a forest population model with demographic rates that respond to the environment, we confirm the predictions of Ellenberg’s classical, albeitonlypartially-evidenced, theory on *F. sylvatica*’s predominance in Central European forests. Furthermore, for thefirsttime, we substantiate the theory by elucidating how the environmental variation in species composition is based in demographic processes. This demonstrates that our approach can be utilized to predict distributions of interacting species and to explain the dynamics between species, as influenced by their environment.

## 1 Introduction

Understanding where species occur is one of the central questions in ecology (MacArthur, 1972; Sutherland et al., 2013). To describe the occurrence of species dependent on environmental variables, ecologists have used the Hutchinson niche concept (Hutchinson, 1957) and abundance response curves (Ellenberg, 1952; Whittaker, 1967; Ter Braak & Prentice, 1988). In the quest to go beyond description, ecologists since Maguire (1973) have sought to explain the change in abundance along environmental gradients with the variation of demographic processes (Watkinson, 1985; Schurr et al., 2012; Normand et al., 2014).

A method for explaining changing species abundance by the environmental responses of the underlying demographic processes, is fitting process-explicit species distribution models (SDM; e.g., Higgins et al., 2012; Schurr et al., 2012; Normand et al., 2014; Pironon et al., 2017; Kunstler et al., 2021; Schultz et al., 2022). Process-explicit SDMs received much attention a decade ago, as part of a research agenda for better forecasting range shifts under climate change (Buckley et al., 2010; Pagel & Schurr, 2012; Zurell et al., 2012; Thuiller et al., 2013; Merow et al., 2014). There is, however, little evidence that these dynamic models predict species ranges much better than simple correlative SDMs, and as a consequence they are not widely used (Morin & Thuiller, 2009; Zurell et al., 2016; Briscoe et al., 2019). But in contrast to purely-correlative SDMs, process-explicit SDMs have untapped potential as a tool for understanding why species occur where they do (Dormann et al., 2012). For instance, they can help understand the relative importance of different demographic processes and interactions with other species dependent on the environment (Schurr et al., 2012; Thuiller et al., 2013; Malchow et al., 2022). Moreover, it has been shown for trees, that early life stages are both critical filters for species composition (Grubb, 1977; Young et al., 2005; Lines et al., 2019) and that saplings can respond differently to the environment than adults (Bertrand et al., 2011; Heiland et al., 2022). Therefore, there have been calls for explicit modelling of seedling and sapling recruitment to capture environmental variation of forest dynamics (Price et al., 2001; Kunstler et al., 2009; Hanbury-Brown et al., 2022; König et al., 2022). Here, we introduce a process-explicit SDM that incorporates the environmental variability of demographic processes for both saplings and adult trees, to link a classical ecological theory on *Fagus sylvatica*’s predominance by Ellenberg 1963 to forest demography.

The Ellenberg theory states that the late-successional tree species *Fagussylvatica* L. (commonbeech, hereafter *Fagus*) is the predominant species at mesic soil conditions of submontane Central European forests, because under these conditions it is more competitive than other species (Ellenberg, 1963). Mesic soil conditions, i.e. conditions without extreme chemical properties as indicated by intermediate to high pH (Härdtle et al., 2004) and without shortages or excess of water, are assumed to be physiologically optimal for most tree species, so that their *physiological response* along soil pH and water gradients would be similar among species–were it not for species interactions (Ellenberg, 1952). In particular, *Fagus* is assumed to outcompete other tree species at mesic soil conditions, so that their *ecological optima* are shifted towards more extreme conditions (Ellenberg, 1963; Keddy, 2001). *Fagus*’ competitive demographics compared to other species include better shade-tolerance of seedlings and saplings (Wagner et al., 2010; Käber et al., 2021), which gives rise to a positive feedback mechanism between sapling recruitment and the shading from the overstory, so that at late successional stages *Fagus* predominates at the competitive equilibrium (see also Peters, 1997; Leuschner & Ellenberg, 2017b; Petrovska et al., 2021a; Heiland et al., 2023). However, through environmental variation of forest demography, *Fagus* ceases to be predominant towards the extremes of the soil gradients.

Although the Ellenberg theory is taken as given throughout research to date (e.g., Barna & Bosela, 2015; Mellert et al., 2016; Leuschner, 2020; Cailleret et al., 2020), only some elements of the theory have been confirmed. The predicted pattern of *Fagus*’ predominance under mesic environmental conditions has been supported by numerous descriptions of natural forests (Peters, 1997; Leuschner et al., 2006; Bolte et al., 2007; Axer et al., 2021; but see minor modifications by Leuschner & Ellenberg, 2017b). But, only some elements of the mechanisms postulated by Ellenberg (1963) have yet been evidenced. Specifically, studies have shown physiological constraints on *Fagus* in highly acidic soil conditions (Leuschner et al., 2006; Aertsen et al., 2012), but not in calcareous soils (Ljungström et al., 1990). Also in wet, waterlogged soils, *Fagus* experiences reduced physiological functioning (Dreyer, 1994; Schmull & Thomas, 2000; Scharnweber et al., 2013; Axer et al., 2021), leading to increased adult mortality (Gorzelak et al., 2000). While *Fagus* is moderately drought-tolerant (Niinemets & Valladares, 2006; Rötzer et al., 2017; Leuschner, 2020), extreme drought conditions makes a plings susceptible to embolism (Tomasella et al., 2019), reduce nitrogen uptake (Fotelli et al., 2001; Geßler et al., 2006; Leuschner, 2020), and inhibit adult growth (Ciais et al., 2005). A study on species interactions by Fichtner et al. (2012) highlighted that competition has a more severe effect on adult beech in less productive environments along soil chemistry and water level gradients and Jacobs et al. (2022) showed that the negative competition effect from *Fagus* on *Quercus petraea* (Matt.) Liebl. disappeared in dry conditions. The specific demographic drivers of species turnover along these soil gradients remain however unclear. *Fagus*’ exceptional shade-tolerance has been confirmed by several studies (Niinemets & Valladares, 2006; Petritan et al., 2007; Käber et al., 2021; Petrovska et al., 2021a). In particular, Heiland et al. (2023) demonstrated with a size-structured forest population model, that shade-tolerance in saplings is the key mechanism for *Fagus*’ predominance at the competitive equilibrium. However, it has not been verified with large-scale environmental data whether demographic rates of *Fagus* indeed vary across the environment in a way that leads to its predominance only under mesic environmental conditions. Moreover, the specific variations in demographic processes that drive the species turnover toward the extremes of soil gradients, remain unclear (see also Meier et al., 2011; Maréchaux et al., 2021). Therefore, here we ask, how environmental variation of demographic rates of *Fagus* and other tree species drives *Fagus*’ relative abundance along environmental gradients?

To tackle this question, we used the JAB forest population model (Heilandetal., 2023). This parsimonious dynamic model explicitly includes three size stages including the sapling stage and the competitive interactions between two “species”, *Fagus sylvatica* and all *other* species. We fitted the model using short time series data from the German NFI, using Bayesian inverse calibration of species-specific and environmentally-responsive demographic rates (Hartig et al., 2012). This allows us to infer how rates of *Fagus sylvatica* vary as a function of soil pH and water level (a two-dimensional environmental space, i.e. ecogram, according to Ellenberg, 1963). With the species-specific and environmentally-responsive rates, we simulated competitive equilibria to (1) test the prediction that *Fagus sylvatica* will be the predominant species at mesic environmental conditions. Further, (2) we used a simulation method to test which demographic rates’ variation explain *Fagus*’ change in relative abundance along the environmental gradients.

## 2 Materials and Methods

### 2.1 JAB Model

The JAB model (Heiland et al., 2023) is a simple dynamic forest population model that includes three size stages including the sapling stage. The model concentrates on competition between multiple species and on the species-specific sapling dynamics by assuming a simple two-layer size structure (see also Valladares & Niinemets, 2008; Cordonnier et al., 2019)

The three stages of the JAB model include the juvenile stage J, representing the understory, and the stages A and B, jointly representing the overstory (Fig. S1). The overstory (BA) is divided into the intermediary stage A and the final stage B, allowing conversion between count density in the sapling stage J and basal area as a measure for competition in the final stage B (Biging & Dobbertin, 1992). The overstory (A and B) shades the understory, but is itself not affected by the understory J (Angelini et al., 2015; Cordonnier et al., 2019; De Lombaerde et al., 2019). Hence, the JAB model captures asymmetric competition (Schwinning, 1998) between overstory and understory.

The JAB model incorporates several key demographic processes that are species-specific (here, we call these processes “demographic rates”, including the limiting competition effects; Table 1). In the sapling stage J, these include the competition response to the overstory’s basal area BA (*s*), competition among saplings (*c_J_*), transition resulting from growth and survival to stage A (*g*), external seedling input (*L_p_* = B*_p_ l*, see Suppl. Section A.2), and seedling regeneration from the local overstory (*r*). Transition from stage A to B is represented by the growth and survival rate *h* together with a conversion factor from counts to basal area (*β_uA_*, see Suppl. Section A.1). Basal area growth in the final stage B is captured by the net increment rate *b*. The overstory stages A and B have competition response to BA (*c_A_* and *c_B_*) so that all stages are logistically limited by the overall tree density through different competition parameters (J: *s*, *c_J_* ; A: *c_A_*; and B: *c_B_*). The density dependence applies not only to the stages but also to the growth rates *g*, *h*, and *b* so that in addition to these rates at the absence of competition, we report estimates of the density-dependent growth terms 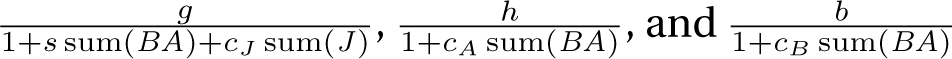 (Table 2 and Fig. S7).

**Table 1:** Demographic rates in the JAB model and their explanation. The rates generally link two model states, so that there is an effect acting from some model state on another that can be positive or negative (indicated in column direction).

| Stage | Symbol | Shorthand | Explanation | Effect from ... | on | (direction) |
| --- | --- | --- | --- | --- | --- | --- |
| J | $L_p/l$ | input | seedling input from large-scale and long-term basal area average to specific J | $\mathfrak{B}$ | J | + |
| | $r$ | recruitment | seedling recruitment from the local basal area to specific J | BA | J | + |
| | $c_J$ | juvenile competition | competition effect from total count of juveniles on specific J | sum J | J | - |
| | $s$ | shading | competition effect from total basal area on specific J | sum BA | J | - |
| J/A | $g$ | transition to A | transition due to survival and growth from specific J to specific A | J/A | A/J | +/- |
| A | $c_A$ | competition on A | competition effect from total basal area on specific A | sum BA | A | - |
| A/B | $h$ | transition to B | survival and growth from specific A to specific B | A/B | B/A | +/- |
| B | $b$ | net basal area growth | net basal area increment of specific B dependent on specific B (also includes mortality) | B | B | + |
| | $c_B$ | competition on B | competition effect from all basal area on specific B (includes both mortality and growth reduction) | sum BA | B | - |

**Table 2:** Posterior distributions of JAB model parameters, including the paraboloid parameters (mean *±* posterior standard deviation) and the environmental variation of log demographic rates (mean *±* environmental standard deviation). Demographic rates are generated by the corresponding paraboloid parameters *ε*, *κ*, *ζ* with the respective environmental axes *x* (pH) and *y* (soil water level). In addition to the growth rates log *g*, log *h* and log *b* the cor-respon_(_ding density-dependent (*d.-d.*) terms across all subpopulations are given at initial and equilibrium state, i.e. 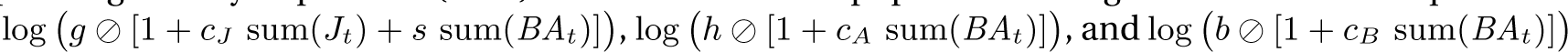.

| Stage | Rate | Parameter | Explanation | Fagus | others | Prior |
| --- | --- | --- | --- | --- | --- | --- |
| J | $r$ | $\log r$ | **env. variation** | **$2.232 \pm 0.1561$** | **$4.629 \pm 0.1366$** | |
| | | $\epsilon_r$ | extremum | $2.235 \pm 0.1560$ | $4.632 \pm 0.1367$ | $4 \pm 1$ |
| | | $\kappa_{r\ x}$ | center (pH) | $0.0001603 \pm 0.9643$ | $0.1004 \pm 0.9937$ | |
| | | $\kappa_{r\ y}$ | center (soil water) | $0.008588 \pm 0.9955$ | $0.01348 \pm 0.9690$ | |
| | | $\zeta_{r\ x}$ | spread (pH) | $-0.0007368 \pm 0.0005630$ | $-0.0007302 \pm 0.0005647$ | |
| | | $\zeta_{r\ y}$ | spread (soil water) | $-0.0007352 \pm 0.0005440$ | $-0.0007166 \pm 0.0005464$ | |
| | $c_J$ | $\log c_J$ | **env. variation** | **$-12.23 \pm 0.2726$** | **$-8.272 \pm 0.7036$** | |
| | | $\epsilon_{c_J}$ | extremum | $-12.79 \pm 0.3719$ | $-8.964 \pm 0.8795$ | $-11 \pm 2$ |
| | | $\kappa_{c_J\ x}$ | center (pH) | $0.4383 \pm 0.6883$ | $0.8527 \pm 0.6355$ | |
| | | $\kappa_{c_J\ y}$ | center (soil water) | $-0.9838 \pm 0.6394$ | $-0.9699 \pm 0.6583$ | |
| | | $\zeta_{c_J\ x}$ | spread (pH) | $0.1329 \pm 0.08853$ | $0.1771 \pm 0.1001$ | |
| | | $\zeta_{c_J\ y}$ | spread (soil water) | $0.1736 \pm 0.09618$ | $0.1731 \pm 0.09071$ | |
| | $s$ | $\log s$ | **env. variation** | **$-9.367 \pm 1.030$** | **$-0.3539 \pm 0.5606$** | |
| | | $\epsilon_s$ | extremum | $-10.25 \pm 1.216$ | $-0.5047 \pm 0.5613$ | $-6 \pm 2$ |
| | | $\kappa_{s\ x}$ | center (pH) | $0.2164 \pm 0.7749$ | $-0.1824 \pm 0.9156$ | |
| | | $\kappa_{s\ y}$ | center (soil water) | $-0.2101 \pm 0.7144$ | $1.039 \pm 0.8310$ | |
| | | $\zeta_{s\ x}$ | spread (pH) | $0.2539 \pm 0.2006$ | $0.02001 \pm 0.01794$ | |
| | | $\zeta_{s\ y}$ | spread (soil water) | $0.3971 \pm 0.3607$ | $0.04323 \pm 0.02879$ | |
| J/A | $g$ | $\log g$ | **env. variation** | **$-5.190 \pm 0.1140$** | **$-2.304 \pm 0.5375$** | |
| | | $\epsilon_g$ | extremum | $-4.863 \pm 0.1734$ | $-2.227 \pm 0.5361$ | $-4 \pm 1$ |
| | | $\kappa_{g\ x}$ | center (pH) | $-0.3004 \pm 0.7416$ | $-0.2853 \pm 0.8740$ | |
| | | $\kappa_{g\ y}$ | center (soil water) | $1.015 \pm 0.5867$ | $0.4063 \pm 0.9424$ | |
| | | $\zeta_{g\ x}$ | spread (pH) | $-0.06200 \pm 0.04883$ | $-0.01870 \pm 0.01612$ | |
| | | $\zeta_{g\ y}$ | spread (soil water) | $-0.1155 \pm 0.05590$ | $-0.02175 \pm 0.01889$ | |
| | | *d.-d. term* | at initial state | $-5.224 \pm 0.1109$ | $-4.895 \pm 0.1050$ | |
| | | | at equilibrium | $-5.254 \pm 0.1090$ | $-6.053 \pm 0.2159$ | |
| A | $c_A$ | $\log c_A$ | **env. variation** | **$-8.859 \pm 1.082$** | **$-6.622 \pm 0.2118$** | |
| | | $\epsilon_{c_A}$ | extremum | $-9.551 \pm 1.328$ | $-6.756 \pm 0.2456$ | $-7 \pm 2$ |
| | | $\kappa_{c_A\ x}$ | center (pH) | $0.1756 \pm 0.7395$ | $0.1406 \pm 0.8022$ | |
| | | $\kappa_{c_A\ y}$ | center (soil water) | $-0.2601 \pm 0.8150$ | $0.1530 \pm 0.8889$ | |
| | | $\zeta_{c_A\ x}$ | spread (pH) | $0.3011 \pm 0.2648$ | $0.04833 \pm 0.04099$ | |
| | | $\zeta_{c_A\ y}$ | spread (soil water) | $0.1924 \pm 0.1923$ | $0.03918 \pm 0.03615$ | |
| A/B | $h$ | $\log h$ | **env. variation** | **$-2.835 \pm 0.1221$** | **$-2.988 \pm 0.07214$** | |
| | | $\epsilon_h$ | extremum | $-2.412 \pm 0.2077$ | $-2.823 \pm 0.08453$ | $-3 \pm 1$ |
| | | $\kappa_{h\ x}$ | center (pH) | $0.9856 \pm 0.5552$ | $-0.09541 \pm 0.8960$ | |
| | | $\kappa_{h\ y}$ | center (soil water) | $0.4131 \pm 0.4774$ | $1.089 \pm 0.4414$ | |
| | | $\zeta_{h\ x}$ | spread (pH) | $-0.1220 \pm 0.06250$ | $-0.01314 \pm 0.01197$ | |
| | | $\zeta_{h\ y}$ | spread (soil water) | $-0.1421 \pm 0.08160$ | $-0.06805 \pm 0.02732$ | |
| | | *d.-d. term* | at initial state | $-2.841 \pm 0.1232$ | $-3.017 \pm 0.07450$ | |
| | | | at equilibrium | $-2.850 \pm 0.1250$ | $-3.062 \pm 0.08182$ | |
| B | $b$ | $\log b$ | **env. variation** | **$-3.546 \pm 1.292$** | **$-3.047 \pm 0.2228$** | |
| | | $\epsilon_b$ | extremum | $-2.947 \pm 0.4956$ | $-3.004 \pm 0.2172$ | $-3 \pm 1$ |
| | | $\kappa_{b\ x}$ | center (pH) | $0.5314 \pm 0.7471$ | $0.006300 \pm 0.9264$ | |
| | | $\kappa_{b\ y}$ | center (soil water) | $-0.4941 \pm 0.9391$ | $0.1035 \pm 0.9095$ | |
| | | $\zeta_{b\ x}$ | spread (pH) | $-0.1114 \pm 0.1490$ | $-0.009622 \pm 0.009471$ | |
| | | $\zeta_{b\ y}$ | spread (soil water) | $-0.1498 \pm 0.2490$ | $-0.01575 \pm 0.01439$ | |
| | | *d.-d. term* | at initial state | $-3.576 \pm 1.281$ | $-3.078 \pm 0.2171$ | |
| | | | at equilibrium | $-3.614 \pm 1.271$ | $-3.126 \pm 0.2115$ | |
| B | $c_B$ | $\log c_B$ | **env. variation** | **$-6.754 \pm 0.6812$** | **$-6.543 \pm 0.1965$** | |
| | | $\epsilon_{c_B}$ | extremum | $-6.916 \pm 0.7710$ | $-6.600 \pm 0.2057$ | $-7 \pm 2$ |
| | | $\kappa_{c_B\ x}$ | center (pH) | $0.1291 \pm 0.8907$ | $0.4474 \pm 0.8884$ | |
| | | $\kappa_{c_B\ y}$ | center (soil water) | $-0.3808 \pm 0.8419$ | $0.4679 \pm 0.8594$ | |
| | | $\zeta_{c_B\ x}$ | spread (pH) | $0.04258 \pm 0.04792$ | $0.01304 \pm 0.01124$ | |
| | | $\zeta_{c_B\ y}$ | spread (soil water) | $0.05732 \pm 0.07456$ | $0.01776 \pm 0.01516$ | |
| (J) | $L_p$ | $\log L_p$ | **env. distribution** | **$2.428 \pm 0.1205$** | **$7.656 \pm 0.1531$** | |
| | | $\log l$ | constant across env. | $1.691 \pm 0.1205$ | $5.095 \pm 0.1531$ | $5 \pm 2$ |

The JAB model can be fitted to short forest time series to infer species-specific demographic rates and their environmental response, extrapolate long-term competitive equilibria (Heiland et al., 2023). As a demographic abundance dynamics model (Briscoe et al., 2019), the model combines the benefits of correlative and process-based approaches, as its parsimony makes it relatively easy to fit to data, yet capable of extrapolating states, assessing the relevance of certain processes, and simulating counterfactual scenarios (Heiland et al., 2023).

For a detailed explanation of the JAB model, see the description in Supporting Information A.1 and Fig. S1, reproduced from Heiland et al. (2023).

### 2.2 Short time series from national forest inventory data

For fitting the JAB model we used short time-series with repeated tree counts and circumference measurements from the German national forest inventory (NFI; main years 1987, 2002, 2012; Table S1).

The forest plots of the German NFI are arranged on a regular quadratic grid with clusters that are 4 km apart, each consisting of four plots that are 150 m apart (Tomppo et al., 2010). In certain federal states, the grid may be intensified to 2.83 km or 2 km between clusters. To fit the JAB model, one randomly selected plot per cluster was used to represent a subpopulation, as the within-cluster variance of tree abundances was high, yet selecting one plot per cluster avoids potential dependencies among the plots of a cluster.

In the German NFI, trees above 10 cm diameter at breast height (dbh) were sampled using the angle count method (Riedel et al., 2017), which selects all trees appearing wider than a defined angle around a central observer at 1.3 m height (Kangas & Maltamo, 2006). Instead of a fixed area for all tree sizes, the selection with an angle leads to a sampling area that varies with dbh. This method delivers data where each counted tree has (1) a corresponding distance to the center, (2) a dbh that is equivalent to a potential sampling distance, i.e. area and sampling probability, (3) as well as a scaling factor *k* [m^2^ ha*^-^*^1^] for the sampling angle. In this study, the angle count plots were truncated to have a fixed maximum radius of 15 m (for details see Section A.3.3).

Trees were grouped into size classes corresponding to the stages of the JAB model. Stages A and B, represented in basal area in the JAB model, included all trees above the lower size threshold for angle count sampling (10 cm). The threshold between A and B was set to dbh 18 cm. Saplings between height 20 cm and dbh 7 cm were counted on circular fixed area plots with radii dependent on size class and survey and were subsumed as size class J in the analyses (Table S2). Because the lower dbh threshold of angle counts, and correspondingly the upper threshold for sapling counts, was changed from 10 cm in 1987 to 7 cm in 2002 in the German NFI, counts for saplings of this size were not available for all NFI years creating a size gap betwen J and A. Despite this gap, in the JAB model fit, we treat the count data in size class J as a proxy for all trees between height 20 cm and dbh 10 cm.

As the analysis focuse son *Fagus sylvatica* we divided trees into two categories (referred to as “species” for brevity): *F. sylvatica* and all *other* species. Within each species and size class, stage abundances were calculated per plot by summing counts in J and A and adding up basal area in A and B. If one of the two species was not present on a plot that was recorded as forested in the NFI, the observation of the respective species was set to 0 for each size class.

To ensure that the NFI data were relevant for Ellenberg’s original predictions related to the environmental gradients and for submontane elevations, we selected plots that that included corresponding records of soil water level (Section 2.3) at elevations between 100 and 600m (these broad altitudinal limits include the submontane level along the different latitudes of the NFI Ellenberg, 1963). Further, to make sure that the data was appropriate for inferring population demographics, we selected repeated observations that were unmanaged during the observation period. Plots were selected that were surveyed thrice—in 1987, 2002, and 2012—which confines the sampled area to former West Germany. Clear cuts or fatal calamities were ruled out by excluding plots that had tree observations in the first or second survey but only zero-observations in a subsequent survey. Plots where categories of management were recorded (sowing, planting, harvested sample trees, or damage through forestry) in the second or third survey were also excluded. Plots with missing sample trees for “unknown reasons” (category only available in the third survey) were also excluded. But unlike Heiland et al. (2023), we did not select the plots based on whether *Fagus* was present or not to sample a greater environmental range of plots.

To reduce computational cost, a random subsample of plots was selected, stratified based on the two environmental variables used in the analysis (Section 2.3). The variables were split into seven uniform intervals between their minimum and maximum values, resulting in 49 bins in environmental space. To achieve an overall sample size below *n* = 1200, either [n/49] = 25 plots were randomly sampled per bin or, in case there were less than 25 plots, all plots were selected (Table S4). Overall, with plot exclusion and sampling criteria, 795 plots were sampled from 61666 plots in 21574 clusters (Table S1).

### 2.3 Environmental variables

We matched the two soil gradients in the original postulation about predominance of *Fagus* (Ellenberg, 1963) by using soil pH and soil water level as environmental predictors in the model fit. For the analyses, we scaled and centered both variables. To visualize forest plots in two-dimensional environmental space, we added a consistent uniform jitter [*-*0.1, 0.1] around the unscaled variables.

#### 2.3.1 Soil pH

We used the top soil pH in CaCl_2_, provided as a spatial raster at 500 m resolution by the European Soil Data Centre (Ballabioetal., 2019). Values of the spatial raster were extracted based on the coordinates of the forest plot locations. Soil pH is commonly used as a proxy for plant nutritional conditions because it is tightly correlated with factors such as base saturation and C/N ratios (Härdtle et al., 2004).

#### 2.3.2 Soil water level

We further used hydromorphic soil water and ground water categories from the National Forest Soil Survey (Benning et al., 2019) at locations of the German NFI to match the soil water gradient of Ellenberg’s prediction. The categories were ranked according to the levels in Ellenberg (1963) and based on expert knowledge and the category descriptions in Benning et al. (2016). To achieve a pseudo-ratio scale, the ordered categories were assigned rational numbers so that the levels between “dry” (−6.0), “damp” (0.0), and “very wet” (8.0) were uniformly spaced (see Suppl. Table S3 for ranking details). The resulting values were averaged per plot, weighted by the respective soil layer thickness and surface area.

#### 2.3.3 Environmental range

To put the environmental range of the German NFI into perspective compared to *Fagus sylvatica*’s European range of distribution, we extracted environmental variables at all plots used in the analysis, and at locations of *Fagus sylvatica*’s presence in Europe (Suppl. Fig. S3). As presences, we used raster cells with predicted relative probability of presence > 1% from The European Atlas of Forest Tree Species (https://forest.jrc.ec.europa.eu/en/european-atlas/; de Rigo et al., 2016). To visualize differences in pH ranges, we used top soil pH in CaCl_2_ (European Soil Data Centre, Section 2.3.1). For comparing ranges of water availability across Europe, instead of water soil level, we used a climatic water balance, described in Heiland et al. (2022).

### 2.4 Fitting the JAB model with environmental response of demographic rates

#### 2.4.1 Model subpopulation structure

In the JAB model, each forest plot is represented as a subpopulation that changes over time. Each sub-population has random initial states of the three stages (J, A, and B), which are fitted to the first observed states of each plot, and then simulated forward. The forward-simulated annual model states are matched to the observed data, with the discrete time *t* after the initial state *t* = 1 in the model to the *t*^th^ year after the first data survey (e.g., 1987). The subpopulation-specific demographic rates (Table 1) are generated dependent on the respective environmental variables (Section 2.4.2).

Using random initial states allowed flexibility for subpopulation trajectories; but to improve convergence and express different levels of certainty we constrained them with gamma priors per species, stage and subpopulation. The gamma distributions were parameterized with mode *µ_y_* and standard deviation *(J’_y_*,

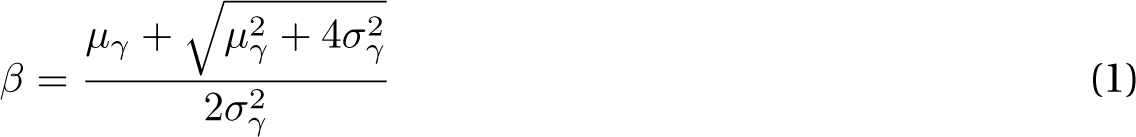

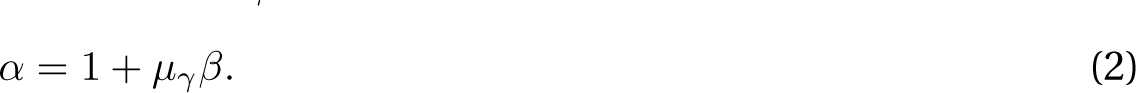

By specifying the mode, we set the highest probability to the observed value, and different standard deviations *(J’* per stage dependent on the observed count *ω* so that *(J’* = [J: 50 + 30*ω*, A: 5+ 3*ω*, B: 0.5+ 0.3*ω*]. Cases with zero observations were parameterized with expected value 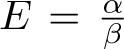 and shape *↵*. By setting *↵* = 1 and *E* to [J: 0.02, A: 0.01, B: 0.005] times the minimum observed value within that stage, we assumed a shape with the maximum probability always near 0 and different degrees of uncertainty for unobserved trees.

#### 2.4.2 Environmental response of demographic rates

Environmental variation of the JAB model rates (*r*, *c_J_*, *s*, *g*, *c_A_*, *h*, *b*, and *c_B_*) is achieved by fitting bell-shaped paraboloids dependent on the two environmental variables *x* (soil pH) and *y* (soil water level). The parabolic response of a demographic rate (*f*) at a subpopulation (*p*) is expressed as the sum of two quadratic response curves,

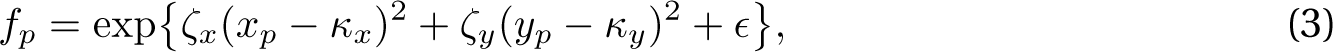

where *ε* corresponds to the optimum of a demographic rate (maximum of the growth-related rates *r*, *g*, *h*, and *b*, and minimum of the competition effects *c_J_*, *s*, *c_A_*, and *c_B_*). The positions of these optima along the two environmental axes *x* and *y* are given by *κ* and the two respective spreads over the axes are given by *ζ*. The exponentiation ensures that all JAB model rates are strictly positive.

This vertex parameterization, with five paraboloid parameters per demographic rate, makes it straight-forward to formulate prior distributions for all parameters. The optimum *ε* determines the general magnitude of the demographic rate and is assumed to be similar to the posterior rate estimates of the JAB model published in Heiland et al. (2023) (for priors see Table 2). The centers *κ* were assumed to conform to a normal distribution *κ ⇠ N* (0, 1). Since the environmental variables (Section 2.3) were scaled and centered, this expresses the belief that the optimum is positioned along the range of the respective variable with a higher probability of intermediate values. The spreads *ζ* determine the direction (bell shape or upside-down bell shape) and the curvature of the paraboloid along the two axes. The growth rates (*r*, *g*, *h*, *b*) were assumed to have negative *ζ*, indicating a maximum at the center and slower growth towards the margins, while the limiting rates (*c_J_*, *s*, *c_A_*, *c_B_*) were assumed to have positive *ζ* with a central minimum and greater limitations towards the margins along both axes. The curvature is assumed to follow a half-normal prior with its greatest mass close to zero, *±ζ ⇠ H*(*(J’_ζ_*), which means that a flat response to the environment with value *ε* is the most likely case a priori. The standard deviation *(J’_ζ_*of the prior was adapted to the prior expected value for *ε* to make it comparable across demographic rates. Comparability is achieved by calculating *(J’_ζ_* = *ζ_x,y_* so that the exponentiated parabola would have a consistent slope *q* = 0.1 at *x, y* = 1 and for *κ* = 0. Given that the spread over the respective other environmental axis *ζ_y_* is 0, the slope *q_x_* corrresponding to a prior expected value of *ε* is *q_x_* = 2*ζ_x_x* exp^(^*ε* + *ζ_x_x*^2)^, and vice versa. Hence, the comparable prior standard deviation for *(J’_ζ_* was calculated as the value of *ζ* where *q* = 0.1, i.e. 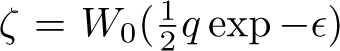, where *W*_0_ is the principal branch of the Lambert *W* function.

All priors were specified equivalently for both species.

In contrast to the other demographic rates, the seedling input rate *l* varies not with the environmental variables, but with a spatial smooth of the regional conspecific basal area, a measure for long-distance and long-term dispersal (details in Suppl. Section A.2). The regional basal area B*_p_* informs the subpopulation-specific values, *L_p_* = exp(log *l*)B*_p_* iterated over the subpopulations or NFI plots *p*.

To characterize the variation of demographic rates along environmental gradients, we took the means of the posterior log rates per subpopulation and summarized this *environmental variation* by calculating means and standard deviations (Table 2).

#### 2.4.3 Likelihood function

The JAB modelstates, including the unknown initial state, were fitted to the data with the same Negative Binomial likelihood function but with varying offsets and observation errors.

Different offsets were used to integrate two kinds of data, both from count processes (see Suppl. Section A.3 for details): (1) Count data in the size classes J and A originate from a count process on varying areas per tree size, so that the area-standardized model state (counts per hectare) is related to the observations with an offset *o*, which is the average area of one sampling plot. (2) Basal area data in size class B also originate from a count process of individual trees with varying areas, but in addition, each tree originally had an individual basal area record. Hence, to relate the model state of *B* (basal area per hectare) to observed counts in size class B, we use a special offset that includes both the average sampling area and a factor 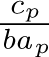, which expresses how many counts *c_p_* are added per unit of basal area *ba_p_* on average per plot. The offsets *o* transform the area-standardized and strictly positive model states (*JAB*) to the counts *C* in the data, so that for each plot *p* within a subpopulation *c*, year *t* (or survey *r*), species *j*, and stage *s*, the likelihood function is expressed by the statement:

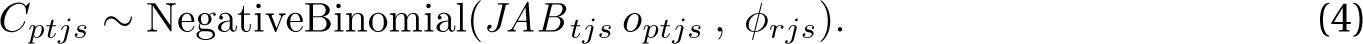

We accounted for variations in sampling protocols and areas by allowing different observation errors, i.e. by fitting the precision parameter *¢* separately for each stage, species, and survey (Table S6). Species-specific *¢* within stage were necessary because the dispersion of a set of multiple species (*others*) is much lower than that of a single species–*F. sylvatica*, which had more stochastic abundance. For the initial state, which was initialized based on the data of the first survey, a different level for *¢* was assumed (Suppl. Table S6). Additionally, changes in sampling areas for size class J between surveys required different values of *¢* for all three surveys in this class. As a result, there were six levels of *¢* in stage J, four levels in stage A, and four levels in stage B. A half-normal prior for the inverse of *¢* was used to improve convergence: 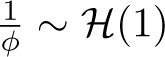.

#### 2.4.4 HMC sampling

The JAB model was implemented in stan (version 2.31.0; Stan Development Team, 2022) and fitted with the package cmdstanr (version 0.5.3; Gabry & Cesnovar, 2021) using its default Hamiltonian Monte Carlo (HMC) algorithm. We used 800 iterations for the warmup and 1000 iterations for the sampling phase, in four independent HMC chains, so that there were 4000 samples of the posterior distributions (see Table 1 for bulk effective sample size, Vehtari et al., 2021). Convergence of the four chains was checked with stan’s default diagnostic *R*^^^ *<* 1.05. All data preparation and posterior analyses were performed with R (version 4.2.3; R Core Team, 2021).

### 2.5 Posterior simulations from the fitted JAB model

Using the posterior probability distributions of the demographic rates, we simulated competitive equilibria and the effect of environmentally-responsive demography on relative abundance. The simulations were implemented in the `generated quantities` block within the stan model.

#### 2.5.1 Competitive equilibria

We simulated the JAB model forward from the initial state to obtain trajectories of subpopulations over time until final equilibrium states were reached. The criterion an equilibrium was that the greatest species-specific relative change in basal area (*BA*) in the last time step should not exceed 1‰: max *|*(*BA_t_ - BA_t-_*_1_) *↵ BA_t_| κ* 0.001. This method for finding the equilibrium points of the JAB models is equivalent to the established procedure for numerical solution of fixed-points of a function as iteration rule (Burden & Faires, 1993). The model was simulated forward at least 250 years to ensure that temporary extrema at the beginning of the trajectory were not mistaken as equilibrium (Heiland et al., 2023), and for a maximum of 6000 years. Additionally, we generated counterfactual equilibria where the regenerative rates (*l*, *r*, *b*) of the respective other species were set to 0, to show the potential abundance of a species in the absence of the other species (Fig. 2). Further, to demonstrate the role of the shading parameter *s*, which had been shown to be the most important for predominance at the equilibrium overall (Heiland et al., 2023), we simulated equilibria with *s* switched between species at the subpopulation level (Fig. 2 G).

Equilibrium abundances were visualized by fitting and predicting two-dimensional tensor smoothing splines along the two environmental axes using the `gam` function of the mgcv package (version 1.8; Wood, 2021). The model was fitted to the posterior distribution with a Gaussian and binomial response for species-specific basal area and its fraction of total basal area, respectively. Predictions from the model were visualized only within the convex hull that included the non-jittered positions of the observed plots, to avoid extrapolation. The line indicating the range of *Fagus* predominance was visualized by predicting the isocline of 50% relative basal area at equilibria (Fig. 1 and Fig. 2 B).

**Figure 1:**
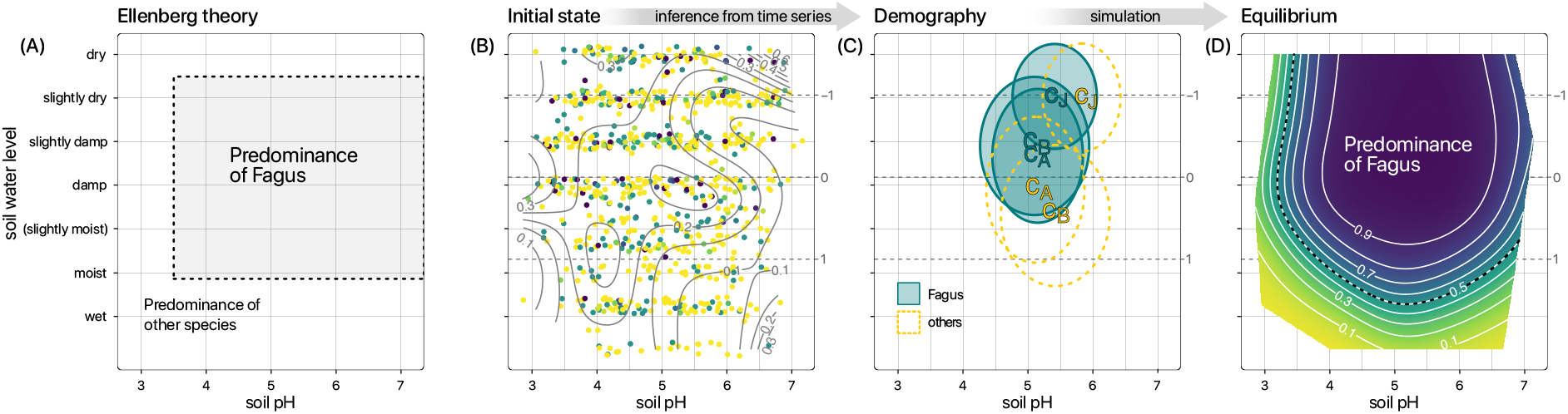
The environmental range along pH and soil water level where *Fagus sylvatica* predominates, as postulated by Ellenberg (1963) (**A**), and the fraction of *Fagus* of the total basal area at the initial subpopulation states, corresponding to the forest plots of the first NFI survey (**B**). As an example for the environmentally-dependent demographic rates (*r*, *cJ*, *s*, *g*, *cA*, *h*, *b*, *cB*), the competition effects *cJ*, *cA*, and *cB* experienced by the size stages J, A, and B are represented here with their minima, i.e. the optima for the abundance of the respective stage (**C**). These optima of the competition effects are visualized by ellipses, representing the standard deviations around the means of the respective center parameters *κ*. The demographic rates, which were inferred from short time series, were extrapolated to yield environmentally-varying competitive equilibria (i.e. fraction of *Fagus* in **D**). Points representing forest plots in environmental space were jittered by 0.1 in both directions. Translation of the hydromorphic soil water levels to English according to Leuschner & Ellenberg, 2017a.

**Figure 2:**
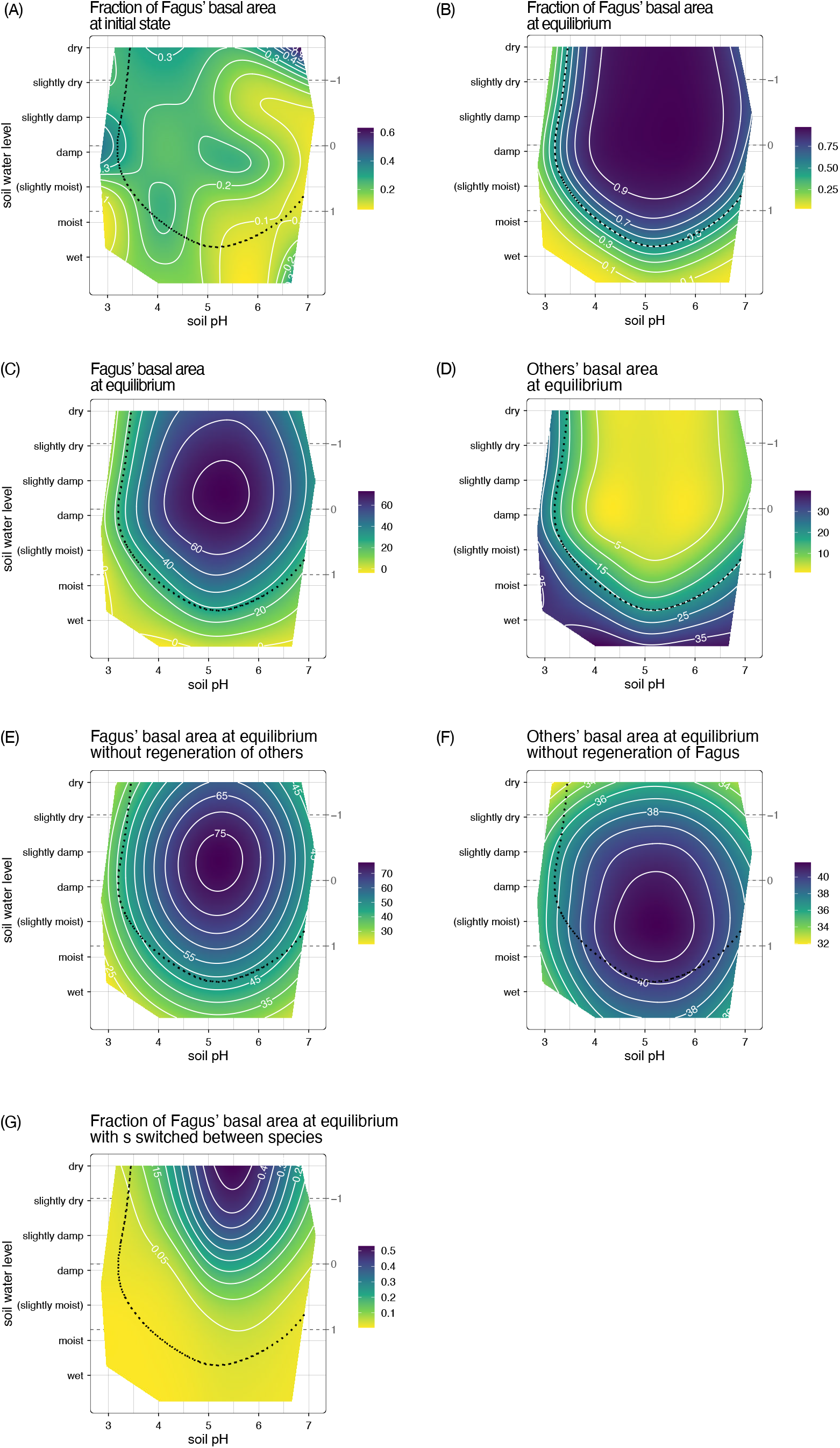
Basal areas in environmental space at initial state (A), at the competitive equilibrium (B–D). Additionally, counterfactual equilibria are depicted: given that the respective other species does not reproduce (by setting *l*, *r*, and *b* to 0; E–F), and (G) given that the shading response (*s*) is switched between species. The contours show predictions from smoothing splines fitted to the posterior distributions (as described in Section 2.5.1)

#### 2.5.2 Effect of environmentally-responsive demography on relative abundance at equilibrium

To quantify the effect of environmentally-responsive demography on the competitive equilibria, we calculated the differences in *Fagus*’ relative basal area at the equilibria with and without environmental variation of the different demographic rates. To simulate counterfactual equilibria without environmental variation of a certain rate, we used the mean rate across subpopulations for simulating all subpopulations. By calculating the differences of the basal area fraction that *Fagus* has at equilibria with flat mean rates and the fraction at equilibria with environmentally-responsive rates, we obtained posterior distributions of effects that environmentally-responsive rates have on *Fagus*’ relative abundance. Negative differences indicate that environmentally-responsive demographic rates have a negative effect on *Fagus*’ basal area at equilibrium. In Fig. 4, we provide a summary of the *elfect of environmentally-responsive demography* for each demographic rate, represented by the mean of the absolute differences.

We further visualized the effect of environmentally-responsive rates in environmental space using Gaussian smoothing splines, as outlined in Section 2.5.1 (Fig. 5 and Suppl. Figs S8). To identify credibly negative or positive effects, we tested if the 90% credible interval intercepted zero or not.

## 3 Results

### 3.1 Equilibria in environmental space

At the equilibria simulated with inferred environmentally-responsive demographic rates, *Fagus* had its highest abundance at the central region of the environmental space. The species predominated at these mesic environmental conditions (basal area fraction *>*50%; Fig. 1, Fig. 2). Even though, at the initial state, *Fagus* had only 14% of the basal are on average (Table S7) with the highest fractions of *Fagus* found at marginal regions (Fig. 2 A). Equilibrium basal area of *Fagus* decreased towards the margins, especially at the wet and at the acidic margin, resulting in *other* species having higher relative basal area. However, at the dry margin, the relative equilibrium basal area of *Fagus* showed minimal decline.

In counterfactual equilibria where regeneration of *others* was set to zero, *Fagus* remained most abundant in the central region of the environmental space. However, in the absence of *Fagus* regeneration, *others* no longer exhibited the highest abundance at the margins but instead showed similar abundance at the central region previously occupied by *Fagus*. Simulating equilibria while switching the shading response (*s*) between species, revealed that without the high shade-tolerance of its saplings *Fagus* would only locally gain a maximum of 50% of the basal area, and it would never become the pre-dominant species (Fig. 2 G).

### 3.2 Environmental variation of demographic rates

In addition to the species differences in mean demographic rates (Suppl. Section B.1), *Fagus* showed much greater environmental variation of its demographic rates compared to *others*. Especially, the demographic rates at the overstory stages A and B had a greater environmental standard deviation on the log scale (environmental variation in Table 2, resulting paraboloids in Fig. 3). At the sapling stage J, however, the environmental variation was more similar between species.

**Figure 3:**
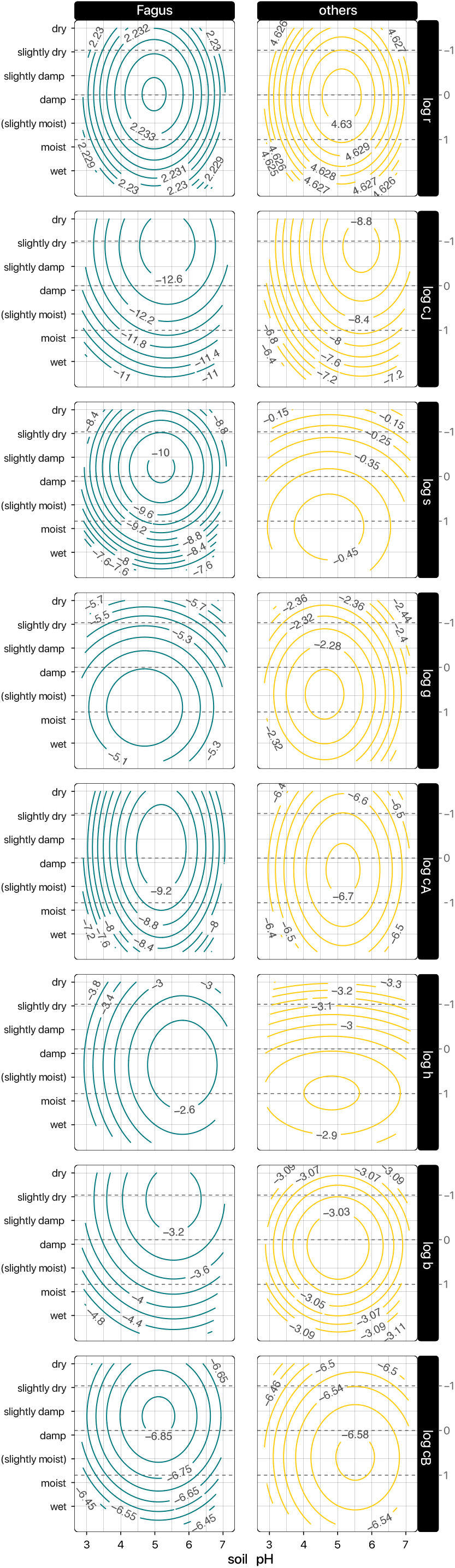
JAB model demographic rates in environmental space. The contours show the mean predicted values from the posterior distribution of polynomials (Section 2.4.2).

The most significant variation overall was observed in the demographic rate log *b* of *Fagus*, which includes both basal area growth and density-independent mortality in overstory stage B, ranging from an optimum of *-*3.5 (Table 2) to values below *-*4.8 (Fig. 3). The competition response of *Fagus* in stage B (log *c_B_*) also varied from a minimum of *-*6.75 to values below *-*6.45. In the smaller size stages, the limiting rates log *c_A_* (stage A), and log *s*, log *c_J_*(stage J) also showed higher variation than the population growth rates, with standard deviations up to 1 on the log scale.

For both species and most demographic rates, the optimum was positioned near the center of both the soil pH and the water level gradient (Fig. 3). Along pH, the optimum for all demographic rates was located close to the center, around moderately acidic values of pH 4.5–6, with a slight tendency towards less acidic conditions indicated by positive *κ_x_* (Table 2). In contrast, along the soil water gradient, there was more variation in the position of the extrema across species, as reflected by larger absolute values of *κ_y_*.

The density-dependent growth terms (2.1) exhibited variations in environmental space similar to the corresponding growth rates (*h* and *b*). The density-dependent transition term for saplings 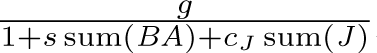 however, had a minimum at the center under equilibrium conditions, reflecting the pattern of the two limiting rates *s* and *c_J_*together with the equilibrium abundance, instead of *g* (Suppl. Fig. S7).

### 3.3 Effect of environmentally-responsive demography on *Fagus*’ relative abundance

The environmental response of *Fagus*’ demographic rates had a much larger effect on relative abundance compared to *others*, i.e. greater absolute mean difference in *Fagus*’ relative basal area at equilibria with environmentally-responsive demographic rates compared to equilibria with flat mean rates (Fig. 4). The most pronounced effects on relative abundance were observed in *Fagus*’ overstory stage B, specifically in the net basal area increment *b* and the limiting rate *c_B_*. The combined environmental response of these two rates also had the largest overall effect on relative abundance. The second-largest effect was observed at the sapling stage J, in the rate *c_J_* . The combination of environmental responses of *g* and of the counteracting rates *c_J_* and *s* also had a considerable effect on relative basal area. In contrast, at the intermediate size stage A, the environmental responses of growth (*h*) and competition (*c_A_*) were not as influential. Interestingly, for *Fagus*, both the seedling recruitment (*r*) and seedling input rate (*L_p_*) exhibited minimal effects on the environmental variation of relative abundance at equilibria.

**Figure 4:**
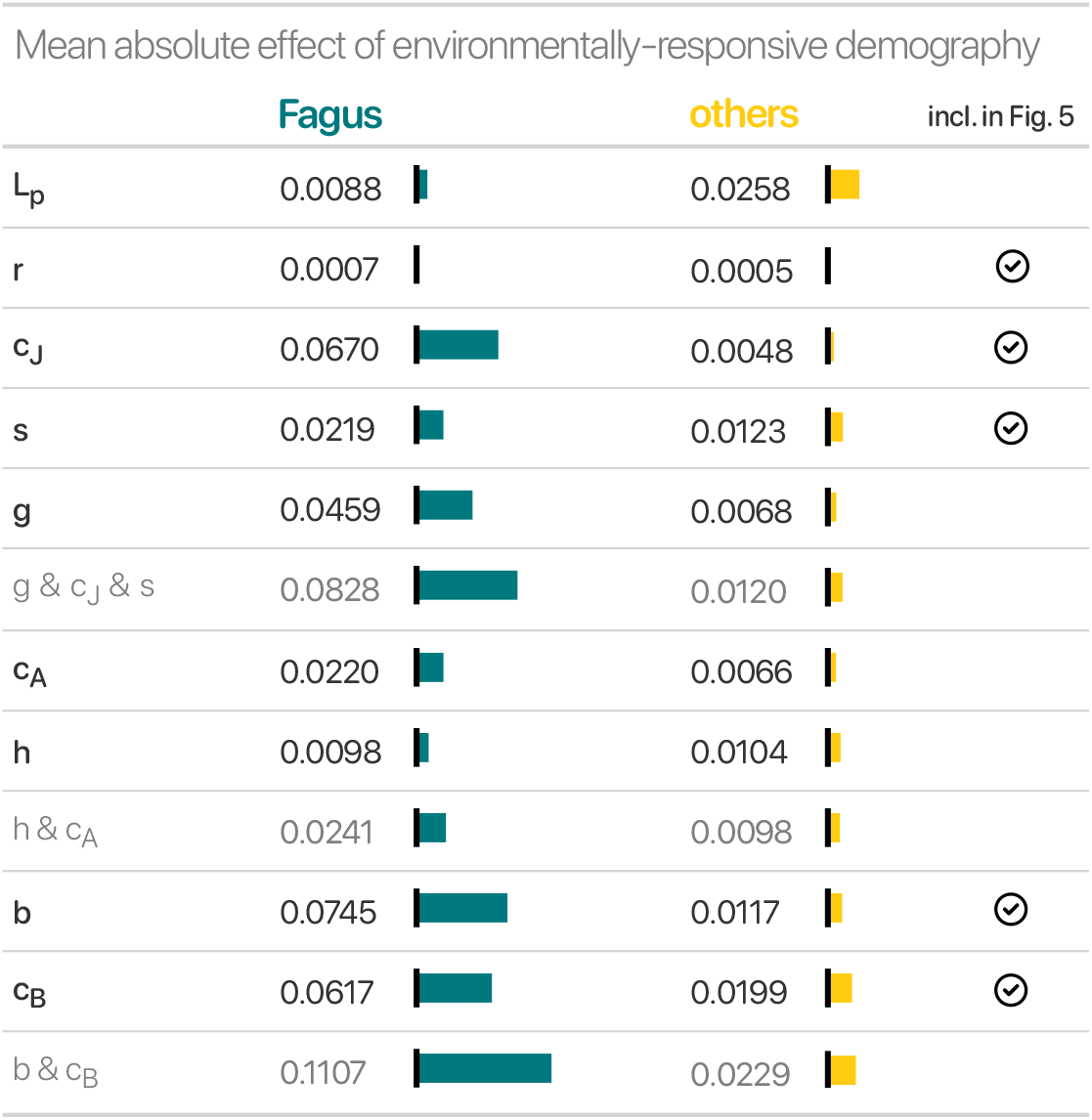
Magnitude of the environmental variation effect per demographic rate (black) and the combined effects of rates in density-dependent terms (grey). The magnitude is indicated by the mean of the absolute effects.

The environmental responses of the demographic rates in *others* had considerably smaller effects on abundance compared to *Fagus*. Although their effect was smaller, in the overstory the relative ranking of the rates’ effects closely resembled that of *Fagus*. Specifically, the response of the rate *c_B_* had the most significant effect on relative abundance, followed by *b*. Among the rates of *others*, the environmental variation of seedling input rate *L_p_* and the shading parameter *s* had the greatest effect on relative abundance at equilibrium. Interestingly, across species, the rate with the smallest effect of the environmental response was seedling recruitment *r*.

The environmental patterns of the three rates with the most significant effects of their environmental response — *b*, *c_J_*, and *c_B_* in *Fagus* — explain the pattern of *Fagus*’ decreasing basal area at the margins (Section 3.2). The environmental response of these rates had a positive effect on *Fagus*’ relative basal area in a central region within its range of predominance, while the range of credibly positive effects is narrower within (90% of the posterior differences *>* 0). Outside the range of predominance, towards the margins, removing the environmental response of these rates negatively affected *Fagus*’ relative abundance, although this negative effect is only credible for a few subpopulations at the very edges (see Fig. 5). In contrast, and deviating from the simulated environmental pattern of relative abundance, the environmental response of the sapling transition rate *g* in *Fagus* (Suppl. Fig. S8), did not result in lower relative basal area under moist conditions. Instead, the response of *g* exhibited a strong positive effect under moist and more acidic conditions whereas a negative influence of declining *g* was observed towards dry conditions. The effect of the environmental variation of the seedling input *L_p_*, despite being the most important within *others*, showed no systematic environmental response (Suppl. Fig. S10).

**Figure 5:**
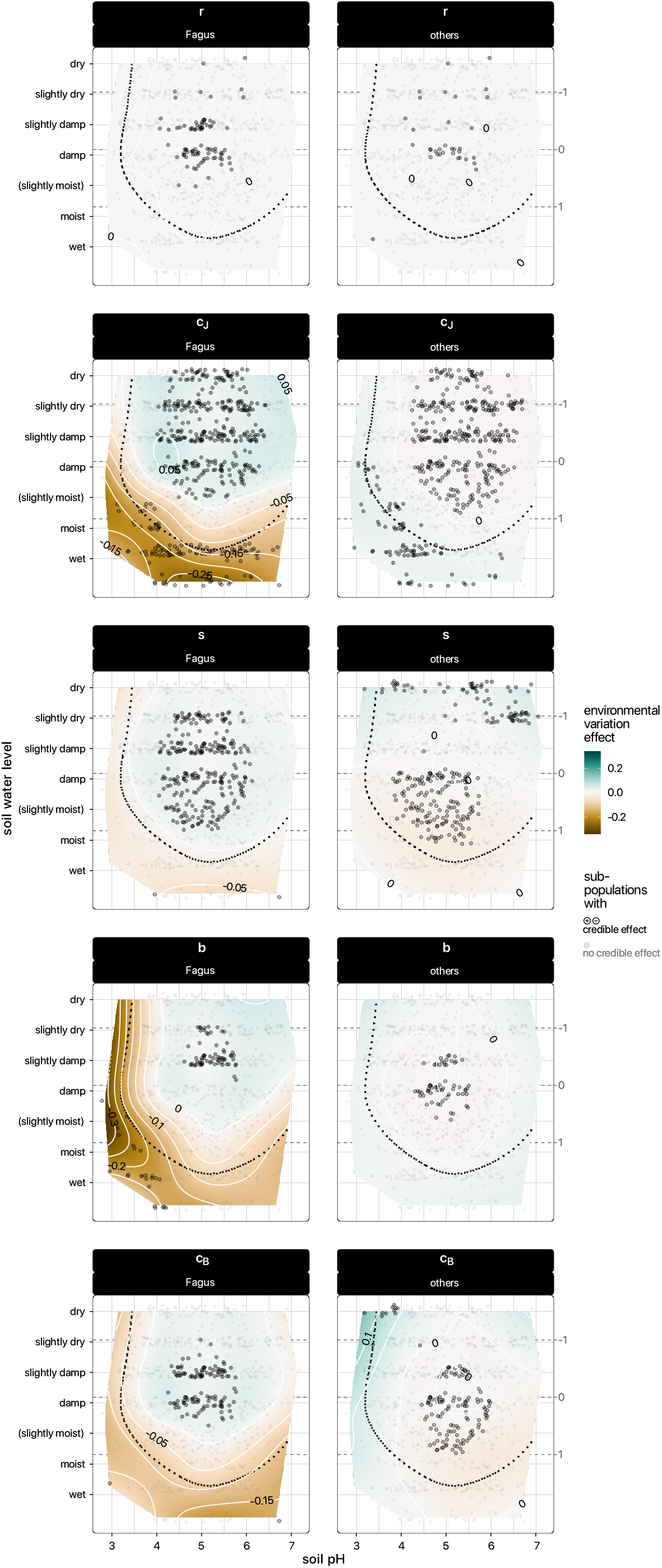
The effect that the environmental response of demographic rates (*r*, *cJ*, *s b*, and *cB*) has on *Fagus*’ relative abundance at equilibrium. The effect of a rate is quantified as the difference between the relative abundances of *Fagus* with and without environmental response of thatrate (Section 2.5.2). Dark point characters indicate credible effects. Effects are assumed as credible if the difference is either *>* or *<* 0 with a probability of *>* 90%. The dashed line indicates the range of *Fagus*’ predominance.

For both species, joint effects of multiple environmentally-responsive rates had a amplified pattern compared to the individual rates alone, for the combination of sapling rates *g*, *c_J_*, and *s*, combined rates of the intermediate stage *h* and *c_A_*, as well as the combined rates of the final stage *b* and *c_B_*. Specifically, the combined effects had a greater positive and negative magnitude, as well as higher credibility compared to their individual effects (Suppl. Fig. S9).

For the environmental patterns of the response effects of all rates see (Suppl. Fig. S8).

## 4 Discussion

In this study, we aimed to test the predictions of a classical ecological theory on species composition in Central European forests (Ellenberg, 1963) by Bayesian inverse calibration of a size-structured forest population model, including two “species”, i.e. *Fagus sylvatica* and *others*. Our simulations largely support the prediction that *Fagus* predominates in mesic environmental conditions while *other* species become more prevalent at the extremes of soil pH and water gradients (Section 4.1). Moreover, we substantiate the theory with demographic analysis (Section 4.2): In a previous study, Heiland et al. (2023) showed that differences in mean demographic rates between *Fagus* and *others*, especially the shade-tolerance of its saplings, explain *Fagus*’ predominance in its core range (these results are largely consistent with the estimates presented in Table 2). Expanding on this approach, here we introduce a model with environmentally-responsive demographic rates to explain the environmental variation of *Fagus*’ relative abundance and why *other* species prevail towards more stressful environmental conditions. Specifically, we show that the environmental variation of overstory demography is greater than that in saplings. As a consequence, we are able to show with a simulation method that the environmental response of adults’ rather than saplings’ demographic rates explains the variation of *Fagus* relative abundance along environmental gradients.

### 4.1 Are simulated equilibrium abundances along environmental gradients consistent with theoretical predictions?

#### 4.1.1 *Fagus* predominates at the center of environmental space

Our findings provide evidence for Ellenberg’s original theory on predominance of *Fagus*. At the competitive equilibrium, *Fagus* is simulated to be the predominant tree species under mesic conditions in a central to dry region of the environmental space while declining towards the margins, as inferred from environmentally-responsive demographic rates with short time series from the German NFI (Section 3.1; Fig. S3). Towards the margins of environmental space, our simulations deviate only slightly from the predictions of Ellenberg (1963). The original prediction postulates a central range of predominance for *Fagus*, but suggests species turnover at the dry margins (Fig. 1). Our results, however, only show minimal decline of *Fagus*’ abundance under dry conditions. This discrepancy may be attributed to three limitations of the data at the dry edge: First, in contrast to Ellenberg’s formulation, the scale available to this study only has the level “dry” and does not differentiate “very dry” (Benning et al., 2019; Suppl. Table S3; Ellenberg, 1963). The lacking differentiation may also have resulted in the much higher number of sampled forest plots at the dry compared to the wet edge (Suppl. Table S4). Second, the sampled data range covers a relatively moist portion of *Fagus*’ range, as evident from the distribution of climatic water balance of the German National Forest Inventory plots compared to the broader range in the European Atlas of Forest Tree Species (means and standard deviations: 215.8 *±* 193.5 vs. 158.3 *±* 343.9 mm a*^-^*^1^; Suppl. Fig. S3; de Rigo et al., 2016). Third, there may be a selection bias in the cultural landscape where extremely dry areas that could be forested (typically with predominance by *Quercus* spp.), have consistently been turned into grasslands throughout Germany (Poschlod, 2015). Consequently, such areas may be excluded from the forest sample, although they have influenced Ellenberg’s theory about natural forests.

In general, both species have a broader distribution along the soil water gradient (Fig. 2), which is at least partially caused by the broader distribution of rate optima along soil water compared to pH (Table 2). The joint effect of broadly-distributed optima rather than steeper curvature of the demographic rates results in a more gradual decline towards the margins and a broader abundance along this gradient (Sharma et al., 2022).

The simulated total equilibrium basal area of around 57 m^2^ ha*^-^*^1^ (Table S7) is only slightly higher than expected from observations. Stillhard et al., 2022 report total basal areas of *⇠*36 m^2^ ha*^-^*^1^ from pristine beech-dominated forests in Uholka-Shyrokyi Luh, Ukraine, while Moreno et al. (2018) project an upper limit of European forests around 50 m^2^ ha*^-^*^1^ (see also Heiland et al., 2023 where the JAB model calibrated with a different selection of plots projected maximum carrying capacities below 50 m^2^ ha*^-^*^1^). Nevertheless, considering that our model does not account for disturbance events like fires, storms, and pathogen damage, the extrapolated equilibria fall within a realistic range and can serve as external validation of the model (e.g., Cailleret et al., 2020).

#### 4.1.2 In *Fagus*’ absence, *others* would also be most abundant at the center of environmental space

In a counterfactual simulation where *Fagus* abundance was minimized by preventing its regeneration, *others* showed maximum abundance in the central region otherwise occupied by *Fagus* (Fig. 2 F). This demonstrates that *others* also have their physiological optimum at the center of the environmental space but are outcompeted by *Fagus*. This mechanism, whereby most species have their physiological optimum under intermediate conditions but species interactions drive the species turnover along environmental gradients (ecological optimum), is a central principle of Ellenberg’s original theory (Ellenberg, 1952, 1963; see also Hutchinson, 1957; Keddy, 2001). The theory also aligns with the stress-gradient hypothesis (Bertness & Callaway, 1994; Käber et al., 2023), which proposes that as stress decreases in favorable conditions competition intensifies, as supported by empirical findings on *Fagus* by Meier et al. (2011). Consequently, it can be concluded that *Fagus* predominates because its demography is disproportionally more beneficial at the center so that it outcompetes *others* (see also Jacobs et al., 2022).

### 4.2 How does the environmental response of demographic rates control relative abundance?

#### 4.2.1 Environmental variation of *Fagus*’ relative abundance is driven by the environmental response of *Fagus*’ demographic rates rather than *others*

Demographic rates of *Fagus* vary more strongly along the environmental gradients compared to *others* (Section 3.2). As a consequence, *Fagus*’ relative abundance and therefore predominance in mesic environmental conditions is primarily driven by the environmental response of *Fagus*’ demographic rates rather than the response of the aggregated *other* species (Fig. 4, Section 3.3). That *Fagus*’ own demographic rates drive its variation in abundance aligns with expectations for a dominant species.

Although the environmental variation of the log demographic rates (Table 2) already hints at the impact on the environmental variation of *Fagus*’ relative abundance, the actual effect that the demographic rates have on equilibrium abundance is also dependent on the interaction with other processes in the JAB model. We therefore quantified environmental effects by comparing equilibria with the given environmentally-responsive demographic rates to equilibria with flat mean demographic rates (Section 2.5.2, Fig. 5). *Fagus*’ demographic rates included the three most important effects of environmentally-dependent demography on relative abundance overall, i.e. *b*, *c_J_*, and *c_B_* (Figs 4 and 5). Among these, environmental response of the net basal area increment *b* had an exceptionally negative effect on *Fagus*’ relative abundance at the acidic edge. This finding aligns with a study by Aertsen et al. (2012), which reports a sharp decrease in *Fagus*’ adult growth when the soil pH drops below 3.6, likely due to aluminum toxicity. In turn, the effect of *b*, which includes both basal area growth and density-independent mortality, does not lead to significant decrease of abundance at the dry edge. This could be attributed to *Fagus*’ moderate drought tolerance (Rötzer et al., 2017; Leuschner, 2020), particularly compared to the most common *other* species, *Picea abies* (L.) H. Karst. (Niinemets & Valladares, 2006; Rötzer et al., 2017; Pretzsch et al., 2020; see Suppl. Table S5 for species composition). Various studies have, however, demonstrated increased mortality under severe drought (Ciais et al., 2005; Geßler et al., 2006; Leuschner, 2020), especially at the southern margin of *Fagus*’ distribution (Jump et al., 2006; Archambeau et al., 2020, here density-independent mortality). In light of this, the insignificant effects of environmentally-responsive demography observed at the dry margin could also be due to the limitations of the soil water scale discussed in Section 4.1. Towards wet conditions, the negative effect of *Fagus*’ environmentally-responsive *b* becomes more pronounced, which aligns with reports about *Fagus* growth sensitivity to waterlogging (Dreyer, 1994; Geßler et al., 2006; Ferner et al., 2012; Durrant et al., 2016 though see Scharnweber et al., 2013). Waterlogging can lead to reduced carbon assimilation due to inhibition of aerobic mitochondrial respiration (Ferner et al., 2012), ultimately resulting in increased mortality (Gorzelak et al., 2000; Geßler et al., 2006).

The effect of *Fagus*’ environmentally-responsive *c_B_* on relative abundance, which represents the adult competition response including mortality and growth reduction, follows a similar pattern as *b* with severe negative effects in wetter conditions (see also Fichtner et al., 2012). However, *c_B_*exhibits less severe negative effects at the acidic margin (Fig. 5). The pattern at the dry margin aligns with findings by Jacobs et al. (2022), who also reported no increased competition effect on *Fagus* growth under drier conditions.

The response of a species’ saplings to competition by all saplings (*c_J_*) is an important process capturing mortality in the JAB model, besides the shading response *s*. Unlike shading, *c_J_* is solely dependent on the density within the sapling stage J itself. This may lead to the environmental response of the rate *c_J_* also capturing environmental variation due to density-independent mortality, since the JAB model disregards density-independent mortality at stage J (Section 4.3). Consequently, here we discuss *c_J_*as a proxy for both density-dependent and density-independent mortality — also given that the literature on environmental variation of *Fagus* sapling mortality does not differentiate between the two. In our study, the environmental response of *Fagus*’ *c_J_* showed positive effects on its relative abundance under neutral and dry conditions, but highly negative effects in acidic and wet conditions. In line with this, a study by Wilkens & Wagner (2021) emphasizes the role of soil fertility for increased survival of small beech saplings, whereas very acidic conditions indicate low soil fertility (Härdtle et al., 2004, although Leuschner & Ellenberg, 2017b found *Fagus* establishment in very acidic soils with thick organic layers). Also in agreement with our results at the upper end of the pH gradient, Härdtle et al. (2004) found no major negative effects on *Fagus* saplings under a liming treatment (Ljungström et al., 1990). Regarding drought tolerance, while Fotelli et al. (2003) found that beech seedlings are more tolerant to dry conditions compared to other co-occurring woody species (see also Madsen & Larsen, 1997), Tomasella et al. (2019) highlight their susceptibility to drought due to their inability to recover from high embolism levels. On the other hand, excess of soil ground water decreases the probability of *Fagus* sapling occurrence (Axer et al., 2021), a potential explanation being species-relatively reduced assimilation, having worse adaption of root formation and carbon allocation to foliage under waterlogging than other species (Dreyer, 1994; Schmull & Thomas, 2000).

#### 4.2.2 Environmental response of overstory demographic rates is more important than the environmental response at earlier stages

The environmental response of demographic rates in adult trees (*b* and *c_b_*in stage B) had the most significant impact on the relative abundance of *Fagus*. Hence, *Fagus* decreasing relative abundance in less favorable environments is mainly due to the environmentally-responsive demography of overstory trees rather than earlier stages. Specifically, the effect of combined environmental responses at the overstory stage (*b* combined with *c_B_*) was greater than the combined effect of rates at the sapling stage (*g* with *s* and *c_J_*). This finding may initially seem surprising, as a previous study has shown that the lower response of *Fagus* saplings to shading (*s*) is the primary factor driving the predominance of *Fagus* at the competitive equilibrium (Heiland et al., 2023; see also Petritan et al., 2007; Leuschner & Ellenberg, 2017b; Petrovska et al., 2021b). However, this apparent contradiction can be resolved. While variation in shade-tolerance is not the main driver of the species’ abundance response in environmental space, the mechanism of shade tolerance remains crucial for *Fagus* predominance in the first place. This is supported by simulated equilibria where the shading response (*s*) was switched between species (Fig. 2 G). Furthermore, both seedling input (*L_p_*), and seedling recruitment (*r*) had no notable effect of environmentally-responsive demography on relative abundance (Suppl. Fig. S10 and Fig. 5), which indicates that the environmental variation of seedling recruitment is better explained by variation of a species’ basal area than by the variation of seedling survival. The smaller effect observed in the environmental response of seedling and sapling rates aligns with the theory that the occurrence of a particular life stage is constrained by the presence of earlier life stages due to “demographic” or “ontogenetic dependency” (Heiland et al., 2022; Young et al., 2005; Ramachandran et al., 2023). Therefore, earlier tree life stages often have broader environmental distributions, as shown by Bertrand et al. (2011); Zhu et al. (2014); Mális et al. (2016); Copenhaver-Parry et al. (2020).

#### 4.2.3 Within overstory demography, the environmental response of the net basal area increment is most important, whereas within earlier size stages, competition response parameters are most important

Within overstory stage B, the rate with the strongest effect of environmentally-responsive demography on *Fagus*’ relative abundance was the net basal area increment *b* of *Fagus*. The net increment does not only include basal area growth, but also density-independent mortality (*b*; see Section 4.3). Consequently, we cannot distinguish between the effects that either of these processes have on the variation of *Fagus*’ relative abundance. On the other hand, it is credible that their joint environmental variation has the greatest effect overall (Fig. 4).

Notably, within the sapling (J) and intermediate stages (A), the environmental variation of *Fagus*’ competition responses *c_J_* and *c_A_*, which capture density-dependent mortality, had the greatest effect on relative abundance (Fig. 4). Similarly, at the overstory (B), the competition response *c_B_* had the second-greatest effect. These competition responses have the function of limiting population growth, counteracting the growth rates (*L_p_*, *r*, *g*, *h*, and *b*). Specifically, within the sapling stage (J), the limiting competition responses (*s* and *c_J_*) had a greater effect than the corresponding growth rates (*r*, *L_p_*, and *g*). That competition effects (*s*, *c_J_*, *c_A_*, *c_B_*) respond to the environment in such a way that they are important for the abundance response, can be explained by their inverse relation to the carrying capacity, i.e. the maximum abundance that the respective environment can support. In the JAB model, the different competition effects jointly determine the carrying capacity, not only of their own but also of the other stages due to coupling via the transition rates (*r*, *g*, *h*). The importance of these limiting processes aligns with previous evidence that individual growth or population growth rates, without accounting for survival, are insufficient predictors for tree species abundance along environmental gradients (Thuiller et al., 2014; Bohner & Diez, 2019; Kunstler et al., 2021).

### 4.3 Limitations of the JAB model and outlook

The JAB model offers a key advantage by allowing inverse calibration with short time series of forest states, rather than demanding detailed demographic data on mortality, growth, and regeneration. This advantage, however, comes with two main limitations that could be addressed through direct estimates of informative priors (Heiland et al., 2023; Hartig et al., 2012).

First, the model includes density-independent mortality only in the net increment *b* of stage B, while stages J and A experience only density-dependent mortality (see Heiland et al., 2023). This simplification was necessary, because without providing further demographic information, density-independent mortality would hardly be separable from the seedling input into the system. Abstracting from density-independent mortality might lead to over estimating competition effects by erroneously absorbing variation due to density-independent mortality, or underestimating growth rates in stages J and A compared to a system with density-independent mortality (see also Section 4.2.1). Future studies could enhance the JAB model by additionally including density-independent mortality, through direct estimation of prior rate distributions with appropriate demographic data on mortality, growth and seedling regeneration.

Second, fitting the model solely to time series of states introduces potential equifinality, where multiple parameter configurations can produce the same model states that fit the data (Hartig et al., 2014). Equifinality in the JAB model became evident through correlations in the posterior estimates of specific rates, such as the optima of overstory rates *ε_b_* and *ε_cB_*. These equifinalities would be a problem for interpretation, if the correlated rates would cancel each other out. To confirm that this is not the case, we check all demographic rates that lead to the abundance within one stage in conjunction, such as *g*, *s*, and *c_J_*, as well as *h* and *c_A_*. This involves comparing the estimates with the full density-dependent growth terms, e.g., *g ↵* [1 + *c_J_* sum(*J_t_*) + *s* sum(*BA_t_*)] (Table 2) and jointly averaging parameters to compute the effect of their joint environmental response (Section 4.2, Fig. 4, Supplementary Fig. S9).

Future studies could overcome these equifinalities by directly estimating individual rates and incorporating more informative priors to constrain the model.

### 4.4 Conclusions

Our study largely supports the prediction of a classical theory (Ellenberg, 1963) that *Fagus sylvatica* naturally predominates in Central European forests under mesic environmental conditions due to its advantageous demography. Towards the extremes of the water availability and pH gradient, *Fagus* relative abundance declines. But contrary to Ellenberg’s original predictions, our results do not show that *others* supplant *Fagus* at the dry margin of the soil water gradient, which could be attributed to limitations of the soil water scale used.

We further explained the environmental pattern of *Fagus*’ predominance with the environmental response of demographic rates by simulating their effects on competitive equilibria. This makes our study the first effort at basing the Ellenberg theory in demography. Our demographic analysis revealed that the environmental change in *Fagus* abundance is primarily driven by variation of its own demographic rates, rather than those of *other* species. Furthermore, we identified that the environmental response of demography with the highest impact on relative basal area occurs at *Fagus*’ overstory stage, even though shade-tolerance of *Fagus* at the sapling stage is the primary factor for its predominance in the first place.

Our study presents a significant breakthrough as it demonstrates that inverse calibration of a simple forest population model with a sapling stage can be used to predict credible patterns of equilibrium abundance along environmental gradients. Moreover, this is the first study that provides a data-driven demographic explanation for a widely-assumed theory on *Fagus* abundance and predominance. This highlights the potential of inverse calibration as an alternative to directly estimating demographic rates to understand species turnover along environmental gradients. By embracing our approach, ecological modelling can predict distributions of interacting species and provide deeper insights into the dynamic interactions between species, as influenced by their environment.

## Author contributions

**L.He.**, together with L.Hü. and G.K., conceived the ideas and designed the methodology; L.He. curated the data; L.He., with advice from L.Hü. and G.K., implemented the software and analysed the data; L.He. wrote the first draft of the manuscript. All authors contributed critically to writing and gave final approval for publication.

## Data Availability Statement

Data available from the Dryad Digital Repository at https://doi.org/10.5061/dryad.3ffbg79pv (Heiland et al. 2023). All software to reproduce the study will be made openly available at https://zenodo.org/.

## References

1. Aertsen, W., Kint, V., De Vos, B., Deckers, J., Van Orshoven, J., & Muys, B. Predicting forest site productivity in temperate lowland from forest floor, soil and litterfall characteristics using boosted regression trees. Plant and Soil, 354(1-2), 157–172 (2012). 10.1007/s11104-011-1052-z.

2. Angelini, A., Corona, P., Chianucci, F., & Portoghesi, L. Structural attributes of stand overstory and light under the canopy. Annals of Silvicultural Research, 39(1), 23–31 (2015). 10.12899/asr-993.

3. Archambeau, J., Ruiz-Benito, P., Ratcliffe, S., Fréjaville, T., Changenet, A., Muñoz Castañeda, J. M., Lehtonen, A., Dahlgren, J., Zavala, M. A., & Benito Garzón, M. Similar patterns of background mortality across Europe are mostly driven by drought in European beech and a combination of drought and competition in Scots pine. Agricultural and Forest Meteorology, 280, 107772 (2020). 10.1016/j.agrformet.2019.107772.

4. Axer, M., Martens, S., Schlicht, R., & Wagner, S. Modelling natural regeneration of European beech in Saxony, Germany: Identifying factors influencing the occurrence and density of regeneration. European Journal of Forest Research (2021). 10.1007/s10342-021-01377-w.

5. Ballabio, C., Lugato, E., Fernández-Ugalde, O., Orgiazzi, A., Jones, A., Borrelli, P., Montanarella, L., & Panagos, P. Mapping LUCAS topsoil chemical properties at European scale using Gaussian process regression. Geoderma, 355, 113912 (2019). 10.1016/j.geoderma.2019.113912.

6. Barna, M. & Bosela, M. Tree species diversity change in natural regeneration of a beech forest under different management. Forest Ecology and Management, 342, 93–102 (2015). 10.1016/j.foreco.2015.01.017.

7. Benning, R., Petzold, R., & Danigel, J. BWI 2012 Umweltdatenbank Bodenprofile. Johann Heinrich von Thünen-Institut, DE (2019).

8. Benning, R., Petzold, R., Danigel, J., Gemballa, R., & Andreae, H. Generating characteristic soil profiles for the plots of the National Forest Inventory in Saxony and. *Waldökologie*, Landschaftsforschung und Naturschutz, 16(2016) (2016).

9. Berger, A., Gschwantner, T., & Schadauer, K. The effects of truncating the angle count sampling method on the Austrian National Forest Inventory. Annals of Forest Science, 77(1), 16 (2020). 10.1007/s13595-019-0907-y.

10. Bertness, M. D. & Callaway, R. Positive interactions in communities. Trends in Ecology & Evolution, 9(5), 191–193 (1994). 10.1016/0169-5347(94)90088-4.

11. Bertrand, R., Gégout, J.-C., & Bontemps, J.-D. Niches of temperate tree species converge towards nutrient-richer conditions over ontogeny. Oikos, 120(10), 1479–1488 (2011). 10.1111/j.1600-0706.2011.19582.x.

12. Biging, G. S. & Dobbertin, M. A comparison of distance-dependent competition measures for height and basal area growth of individual conifer trees. Forest Science, 38(3), 695–720 (1992). 10.1093/forestscience/38.3.695.

13. Bohner, T. & Diez, J. Extensive mismatches between species distributions and performance and their relationship to functional traits. *Ecology Letters*, p. ele.13396 (2019). 10.1111/ele.13396.

14. Bolte, A., Czajkowski, T., & Kompa, T. The north-eastern distribution range of European beech a review. Forestry, 80(4), 413–429 (2007). 10.1093/forestry/cpm028.

15. Briscoe, N. J., Elith, J., Salguero-Gómez, R., Lahoz-Monfort, J. J., Camac, J. S., Giljohann, K. M., Holden, M. H., Hradsky, B. A., Kearney, M. R., McMahon, S. M., Phillips, B. L., Regan, T. J., Rhodes, J. R., Vesk, P. A., Wintle, B. A., Yen, J. D. L., & Guillera-Arroita, G. Forecasting species range dynamics with process-explicit models: Matching methods to applications. Ecology Letters, 22(11), 1940–1956 (2019). 10.1111/ele.13348.

16. Buckley, L. B., Urban, M. C., Angilletta, M. J., Crozier, L. G., Rissler, L. J., & Sears, M. W. Can mechanism inform species’ distribution models? Ecology Letters, 13(8), 1041–1054 (2010). 10.1111/j.1461-0248.2010.01479.x.

17. Burden, R. & Faires, J. Numerical Analysis. Mathematics Series. PWS-Kent Publishing Company, Whitstable (1993).

18. Cailleret, M., Bircher, N., Hartig, F., Bugmann, H., & Hu, L. Bayesian calibration of a growth-dependent tree mortality model to simulate the dynamics of European temperate forests. Ecological Applications, 30(1), 17 (2020). 10.1002/eap.2021.

19. Ciais, Ph., Reichstein, M., Viovy, N., Granier, A., Ogée, J., Allard, V., Aubinet, M., Buchmann, N., Bernhofer, Chr., Carrara, A., Chevallier, F., De Noblet, N., Friend, A. D., Friedlingstein, P., Grünwald, T., Heinesch, B., Keronen, P., Knohl, A., Krinner, G., Loustau, D., et al. Europe-wide reduction in primary productivity caused by the heat and drought in 2003. Nature, 437(7058), 529–533 (2005). 10.1038/nature03972.

20. Copenhaver-Parry, P. E., Carroll, C. J. W., Martin, P. H., & Talluto, M. V. Multi-scale integration of tree recruitment and range dynamics in a changing climate. Global Ecology and Biogeography, 29(1), 102–116 (2020). 10.1111/geb.13012.

21. Cordonnier, T., Smadi, C., Kunstler, G., & Courbaud, B. Asymmetric competition, ontogenetic growth and size inequality drive the difference in productivity between two-strata and one-stratum forest stands. Theoretical Population Biology, 130, 83–93 (2019). 10.1016/j.tpb.2019.07.001.

22. De Lombaerde, E., Verheyen, K., Van Calster, H., & Baeten, L. Tree regeneration responds more to shade casting by the overstorey and competition in the understorey than to abundance per se. Forest Ecology and Management, 450, 117492 (2019). 10.1016/j.foreco.2019.117492.

23. de Rigo, D., Caudullo, G., Houston Durrant, T., & San-Miguel-Ayanz, J. The European Atlas of Forest Tree Species: Modelling, Data and Information on Forest Tree Species, pp. e01aa69+. Publications Office of the European Union, Luxembourg (2016). 10.2760/776635.

24. Din, Q. Dynamics of a discrete Lotka-Volterra model. Advances in Dilference Equations, 2013(95), 1–13 (2013). 10.1186/1687-1847-2013-95.

25. Dormann, C. F., Schymanski, S. J., Cabral, J., Chuine, I., Graham, C., Hartig, F., Kearney, M., Morin, X., Römermann, C., Schröder, B., & Singer, A. Correlation and process in species distribution models: Bridging a dichotomy. Journal of Biogeography, 39(12), 2119–2131 (2012). 10.1111/j.1365-2699.2011.02659.x.

26. Dreyer, E. Compared sensitivity of seedlings from 3 woody species (Quercus robur L, Quercus rubra L and Fagus silvatica L) to water-logging and associated root hypoxia: Effects on water relations and photosynthesis. Annales des Sciences Forestières, 51(4), 417–428 (1994). 10.1051/forest:19940407.

27. Durrant, T. H., de Rigo, D., & Caudullo, G. Fagus sylvatica in Europe: Distribution, habitat, usage and threats. In J. San-Miguel-Ayanz, D. de Rigo, G. Caudullo, T. Houston Durrant, & A. Mauri (editors), European Atlas of Forest Tree Species, pp. e012b90+. Publ. Off. EU,, Luxembourg (2016).

28. Ellenberg, H. Physiologisches und ökologisches Verhalten derselben Pflanzenarten. Berichte der Deutschen Botanischen Gesellschaft, 65(10), 350–361 (1952). 10.1111/j.1438-8677.1953.tb00671.x.

29. Ellenberg, H. Vegetation Mitteleuropas Mit Den Alpen. In Kausaler, Dynamischer Und Historischer Sicht., volume IV/2 of Einführung in Die Phytologie. Ulmer, Stuttgart, 1 edition (1963).

30. Ellner, S. Asymptotic behavior of some stochastic difference equation population models. Journal Of Mathematical Biology, 19(2), 169–200 (1984). 10.1007/BF00277745.

31. Ferner, E., Rennenberg, H., & Kreuzwieser, J. Effectof floodingon C metabolismof flood-tolerant(Quercus robur) and non-tolerant (Fagus sylvatica) tree species. Tree Physiology, 32(2), 135–145 (2012). 10.1093/treephys/tps009.

32. Fichtner, A., Sturm, K., Rickert, C., Härdtle, W., & Schrautzer, J. Competition response of European beech *Fagus sylvatica* L. varies with tree size and abiotic stress: Minimizing anthropogenic disturbances in forests. Journal of Applied Ecology, 49(6), 1306–1315 (2012). 10.1111/j.1365-2664.2012.02196.x.

33. Fotelli, M. N., Geßler, A., Peuke, A. D., & Rennenberg, H. Drought affects the competitive interactions between *Fagus sylvatica* seedlings and an early successional species, *Rubus fruticosus* : Responses of growth, water status and *5* ^13^ C composition. New Phytologist, 151(2), 427–435 (2001). 10.1046/j.1469-8137.2001.00186.x.

34. Fotelli, M. N., Rennenberg, H., Holst, T., Mayer, H., & Geßler, A. Carbon isotope composition of various tissues of beech (*Fagus sylvatica*) regeneration is indicative of recent environmental conditions within the forest understorey. New Phytologist, 159(1), 229–244 (2003). 10.1046/j.1469-8137.2003.00782.x.

35. Gabry, J. & Cesnovar, R. Cmdstanr: R Interface to CmdStan. https://mc-stan.org/cmdstanr/reference/index.html (2021).

36. Gause, G. F. Experimental studies on the struggle for existence: I. Mixed population of two species of yeast. Journal of experimental biology, 9(4), 389–402 (1932). 10.1242/jeb.9.4.389.

37. Geßler, A., Keitel, C., Kreuzwieser, J., Matyssek, R., Seiler, W., & Rennenberg, H. Potential risks for European beech (Fagus sylvatica L.) in a changing climate. Trees, 21(1), 1–11 (2006). 10.1007/s00468-006-0107-x.

38. Goldberg, D. E. Competitive ability: Definitions, contingency and correlated traits. Philosophical Transactionsofthe Royal Society of London. Series B: Biological Sciences, 351(1345), 1377–1385 (1996). 10.1098/rstb.1996.0121.

39. Goldberg, D. E. & Landa, K. Competitive Effect and Response: Hierarchies and Correlated Traits in the Early Stages of Competition. The Journal of Ecology, 79(4), 1013 (1991). 10.2307/2261095.

40. Gorzelak, A. et al. Effect of flooding on the flora – the example of the flooding of the Oder in 1997. Beiträge für Forstwirtschaft und Landschaftsökologie, 34(1), 8–11 (2000).

41. Grubb, P. J. Maintenance of Species-Richness in Plant Communities - Importance of Regeneration Niche. Biological Reviews of the Cambridge Philosophical Society, 52(1), 107–145 (1977). 10.1111/j.1469-185X.1977.tb01347.x.

42. Hanbury-Brown, A. R., Ward, R. E., & Kueppers, L. M. Forest regeneration within Earth system models: Current process representations and ways forward. *New Phytologist*, p. nph.18131 (2022). 10.1111/nph.18131.

43. Härdtle, W., von Oheimb, G., Friedel, A., Meyer, H., & Westphal, C. Relationship between pH-values and nutrient availability in forest soils – the consequences for the use of ecograms in forest ecology. *Flora - Morphology, Distribution*, Functional Ecology of Plants, 199(2), 134–142 (2004). 10.1078/0367-2530-00142.

44. Hartig, F., Dislich, C., Wiegand, T., & Huth, A. Technical Note: Approximate Bayesian parameterization of a process-based tropical forest model. Biogeosciences, 11(4), 1261–1272 (2014). 10.5194/bg-11-1261-2014.

45. Hartig, F., Dyke, J., Hickler, T., Higgins, S. I., O’Hara, R. B., Scheiter, S., & Huth, A. Connecting dynamic vegetation models to data-an inverse perspective. Journalof Biogeography, 39(12), 2240–2252 (2012). 10.1111/j.1365-2699.2012.02745.x.

46. Heiland, L., Kunstler, G., Ruiz-Benito, P., Buras, A., Dahlgren, J., & Hülsmann, L. Divergent occurrences of juvenile and adult trees are explained by both environmental change and ontogenetic effects. Ecography, 2022(3), e06042 (2022). 10.1111/ecog.06042.

47. Heiland, L., Kunstler, G., Seben, V., & Hülsmann, L. Which demographic processes control competitive equilibria? Bayesian calibration of a size-structured forest population model. Ecology and Evolution, 13(7), e10232 (2023). 10.1002/ece3.10232.

48. Higgins, S. I., O’Hara, R. B., Bykova, O., Cramer, M. D., Chuine, I., Gerstner, E.-M., Hickler, T., Morin, X., Kearney, M. R., Midgley, G. F., & Scheiter, S. A physiological analogy of the niche for projecting the potential distribution of plants. Journal of Biogeography, 39(12), 2132–2145 (2012). 10.1111/j.1365-2699.2012.02752.x.

49. Hutchinson, G. E. Concluding Remarks. Cold Spring Harbor Symposia on Quantitative Biology, 22(0), 415–427 (1957). 10.1101/sqb.1957.022.01.039.

50. Jacobs, K., Jonard, M., Muys, B., & Ponette, Q. Shifts in dominance and complementarity between sessileoakandbeechalongecologicalgradients. Journalof Ecology, 110(10), 2404–2417 (2022). 10.1111/1365-2745.13958.

51. Jump, A. S., Hunt, J. M., & Peñuelas, J. Rapid climate change-related growth decline at the southern range edge of *Fagus sylvatica*: *FAGUS* GROWTH DECLINE. Global Change Biology, 12(11), 2163– 2174 (2006). 10.1111/j.1365-2486.2006.01250.x.

52. Käber, Y., Bigler, C., HilleRisLambers, J., Hobi, M., Nagel, T. A., Aakala, T., Blaschke, M., Brang, P., Brzeziecki, B., Carrer, M., Cateau, E., Frank, G., Fraver, S., Idoate-Lacasia, J., Holik, J., Kucbel, S., Leyman, A., Meyer, P., Motta, R., Samonil, P., et al. Sheltered or suppressed? Tree regeneration in unmanaged European forests. Journal of Ecology, 111(10), 2281–2295 (2023). 10.1111/1365-2745.14181.

53. Käber, Y., Meyer, P., Stillhard, J., De Lombaerde, E., Zell, J., Stadelmann, G., Bugmann, H., & Bigler, C. Tree recruitment is determined by stand structure and shade tolerance with uncertain role of climate and water relations. *Ecology and Evolution*, p. ece3.7984 (2021). 10.1002/ece3.7984.

54. Kangas, A. & Maltamo, M. Forest Inventory. Methodology and Applications. Number 10 in Managing Forest Ecosystems. Springer, Dordrecht (2006).

55. Keddy, P. A. Competition. Number 26 in Population and Community Biology Series. Kluwer Academic Publishers, Dordrecht ; Boston, 2nd ed edition (2001).

56. König, L. A., Mohren, F., Schelhaas, M.-J., Bugmann, H., & Nabuurs, G.-J. Tree regeneration in models of forest dynamics – Suitability to assess climate change impacts on European forests. Forest Ecology and Management, 520, 120390 (2022). 10.1016/j.foreco.2022.120390.

57. Kunstler, G., Coomes, D. A., & Canham, C. D. Size-dependence of growth and mortality influence the shade tolerance of trees in a lowland temperate rain forest. Journal of Ecology, 97(4), 685–695 (2009). 10.1111/j.1365-2745.2009.01482.x.

58. Kunstler, G., Guyennon, A., Ratcliffe, S., Rüger, N., Ruiz-Benito, P., Childs, D. Z., Dahlgren, J., Lehtonen, A., Thuiller, W., Wirth, C., Zavala, M. A., & Salguero-Gomez, R. Demographic performance of European tree species at their hot and cold climatic edges. Journal of Ecology, 109(2), 1041–1054 (2021). 10.1111/1365-2745.13533.

59. Leuschner, C. Drought response of European beech (Fagus sylvatica L.)—A review. Perspectives in Plant Ecology, Evolution and Systematics, 47, 125576 (2020). 10.1016/j.ppees.2020.125576.

60. Leuschner, C. & Ellenberg, H. Ecology of Central European Forests: Vegetation Ecology of Central Europe, Volume I. Springer International Publishing, Cham (2017a). 10.1007/978-3-319-43042-3.

61. Leuschner, C. & Ellenberg, H. Ecology of Central European Non-Forest Vegetation: Coastal to Alpine, Natural to Man-Made Habitats: Vegetation Ecology of Central Europe, Volume II. Springer International Publishing, Cham (2017b). 10.1007/978-3-319-43048-5.

62. Leuschner, C., Meier, I. C., & Hertel, D. On the niche breadth of Fagus sylvatica: Soil nutrient status in 50 Central European beech stands on a broad range of bedrock types. Annals of Forest Science, 63(4), 355–368 (2006). 10.1051/forest:2006016.

63. Levine, J. M. & Rees, M. Effects of Temporal Variability on Rare Plant Persistence in Annual Systems. The American Naturalist, 164(3), 350–363 (2004). 10.1086/422859.

64. Lines, E. R., Zavala, M. A., Ruiz-Benito, P., & Coomes, D. A. Capturing juvenile tree dynamics from count data using Approximate Bayesian Computation. *Ecography*, p. ecog.04824 (2019). 10.1111/ecog.04824.

65. Ljungström, M., Gyllin, M., & Nihlgård, B. Effects of liming on soil acidity and beech *(Fagus sylvatica* L.) regeneration on acid soils in south Swedish beech forests. Scandinavian Journal of Forest Research, 5(1-4), 243–254 (1990). 10.1080/02827589009382609.

66. MacArthur, R. H. *Geographical Ecology: Patterns in the Distribution of Species*. Harper & Row, New York (1972).

67. Madsen, P. & Larsen, J. B. Natural regeneration of beech Fagus sylvatica L. with respect to canopy density, soil moisture and soil carbon content. Forest Ecology and Management, 97(2), 95–105 (1997). 10.1016/S0378-1127(97)00091-1.

68. Maguire, B. Niche Response Structure and the Analytical Potentials of Its Relationship to the Habitat. The American Naturalist, 107(954), 213–246 (1973). 10.1086/282827.

69. Malchow, A.-K., Hartig, F., Reeg, J., Kéry, M., & Zurell, D. Demography-environment relationships improve mechanistic understanding of range dynamics under climate change. Preprint, Ecology (2022). 10.1101/2022.09.23.509134.

70. Mális, F., Kopecký, M., Petrík, P., Vladovic, J., Merganic, J., & Vida, T. Life stage, not climate change, explains observed tree range shifts. Global Change Biology, 22(5), 1904–1914 (2016). 10.1111/gcb.13210.

71. Maréchaux, I., Langerwisch, F., Huth, A., Bugmann, H., Morin, X., Reyer, C. P., Seidl, R., Collalti, A., Dantas de Paula, M., Fischer, R., Gutsch, M., Lexer, M. J., Lischke, H., Rammig, A., Rödig, E., Sakschewski, B., Taubert, F., Thonicke, K., Vacchiano, G., & Bohn, F. J. Tackling unresolved questions in forest ecology: The past and future role of simulation models. *Ecology and Evolution*, p. ece3.7391 (2021). 10.1002/ece3.7391.

72. Meier, E. S., Edwards Jr, T. C., Kienast, F., Dobbertin, M., & Zimmermann, N. E. Co-occurrence patterns of trees along macro-climatic gradients and their potential influence on the present and future distribution of Fagus sylvatica L.: Influence of co-occurrence patterns on Fagus sylvatica. Journal of Biogeography, 38(2), 371–382 (2011). 10.1111/j.1365-2699.2010.02405.x.

73. Mellert, K. H., Ewald, J., Hornstein, D., Dorado-Liñán, I., Jantsch, M., Taeger, S., Zang, C., Menzel, A., & Kölling, C. Climatic marginality: A new metric for the susceptibility of tree species to warming exemplified by Fagus sylvatica (L.) and Ellenberg’s quotient. European Journal of Forest Research, 135(1), 137–152 (2016). 10.1007/s10342-015-0924-9.

74. Merow, C., Latimer, A. M., Wilson, A. M., McMahon, S. M., Rebelo, A. G., & Silander, J. A. On using integral projection models to generate demographically driven predictions of species’ distributions: Development and validation using sparse data. Ecography, 37(12), 1167–1183 (2014). 10.1111/ecog.00839.

75. Moreno, A., Neumann, M., & Hasenauer, H. Climate limits on European forest structure across space and time. Global and Planetary Change, 169, 168–178 (2018). 10.1016/j.gloplacha.2018.07.018.

76. Morin, X. & Thuiller, W. Comparing niche- and process-based models to reduce prediction uncertainty in species range shifts under climate change. Ecology, 90(5), 1301–1313 (2009). 10.1890/08-0134.1.

77. Niinemets, Ü. & Valladares, F. Tolerance to Shade, Drought, and Waterlogging of Temperate Northern Hemisphere Trees and Shrubs. Ecological Monographs, 76(4), 521–547 (2006). 10.1890/0012-9615(2006)076[0521:TTSDAW]2.0.CO;2.

78. Normand, S., Zimmermann, N. E., Schurr, F. M., & Lischke, H. Demography as the basis for understanding and predicting range dynamics. Ecography, 37(12), 1149–1154 (2014). 10.1111/ecog.01490.

79. Pagel, J. & Schurr, F. M. Forecasting species ranges by statistical estimation of ecological niches and spatial population dynamics: Statistical estimation of dynamic range models. Global Ecology and Biogeography, 21(2), 293–304 (2012). 10.1111/j.1466-8238.2011.00663.x.

80. Peters, R. Beech Forests. Springer Netherlands, Dordrecht(1997).

81. Petritan, A. M., Von Lupke, B., & Petritan, I. C. Effects of shade on growth and mortality of maple (Acer pseudoplatanus), ash (Fraxinus excelsior) and beech (Fagus sylvatica) saplings. Forestry, 80(4), 397– 412 (2007). 10.1093/forestry/cpm030.

82. Petrovska, R., Brang, P., Gessler, A., Bugmann, H., & Hobi, M. L. Grow slowly, persist, dominate—Explainingbeech dominance in a primeval forest. Ecology and Evolution, p. ece3.7800 (2021a). 10.1002/ece3.7800.

83. Petrovska, R., Bugmann, H., Hobi, M. L., Ghosh, S., & Brang, P. Survival time and mortality rate of regeneration in the deep shade of a primeval beech forest. European Journalof Forest Research (2021b). 10.1007/s10342-021-01427-3.

84. Pironon, S., Villellas, J., Thuiller, W., Eckhart, V., Geber, M., A. Moeller, D., & García, M. The ’Hutchinsonian niche’ as an assemblage of demographic niches: Implications for species geographic ranges. Ecography (2017). 10.1111/ecog.03414.

85. Poschlod, P. Geschichte der Kulturlandschaft: Entstehungsursachen und Steuerungsfaktoren der Entwicklung der Kulturlandschaft, Lebensraum-und Artenvielfalt in Mitteleuropa; 38 Tabellen. Ulmer, Stuttgart (Hohenheim) (2015).

86. Pretzsch, H., Grams, T., Häberle, K. H., Pritsch, K., Bauerle, T., & Rötzer, T. Growth and mortality of Norway spruce and European beech in monospecific and mixed-species stands under natural episodic andexperimentallyextendeddrought. Resultsof the KROOF throughfallexclusionexperiment. Trees, 34(4), 957–970 (2020). 10.1007/s00468-020-01973-0.

87. Price, D. T., Zimmermann, N. E., van der Meer, P. J., Lexer, M. J., Leadley, P., Jorritsma, I. T. M., Schaber, J., Clark, D. F., Lasch, P., McNulty, S., Wu, J., & Smith, B. Regeneration in Gap Models: Priority Issues for Studying Forest Responses to Climate Change. Climatic Change, 51(3), 475–508 (2001). 10.1023/A:1012579107129.

88. R Core Team. R: A Language and Environment for Statistical Computing. R Foundation for Statistical Computing, Vienna, Austria (2021).

89. Ramachandran, A., Huxley, J. D., McFaul, S., Schauer, L., Diez, J., Boone, R., Madsen-Hepp, T., McCann, E., Franklin, J., Logan, D., Rose, M. B., & Spasojevic, M. J. Integrating ontogeny and ontogenetic dependency into community assembly. Journal of Ecology, pp. 1365–2745.14132 (2023). 10.1111/1365-2745.14132.

90. Riedel, T., Hennig, P., Kroiher, F., Polley, H., Schmitz, F., & Schwitzgebel, F. *Die Dritte Bundeswaldinventur (BWI 2012). Inventur-Und Auswertemethoden*. Thünen-Institut, Braunschweig (2017).

91. Rötzer, T., Biber, P., Moser, A., Schäfer, C., & Pretzsch, H. Stem and root diameter growth of European beech and Norway spruce under extreme drought. Forest Ecology and Management, 406, 184–195 (2017). 10.1016/j.foreco.2017.09.070.

92. Scharnweber, T., Manthey, M., & Wilmking, M. Differential radial growth patterns between beech (Fagus sylvatica L.) and oak (Quercus robur L.) on periodically waterlogged soils. Tree Physiology, 33(4), 425–437 (2013). 10.1093/treephys/tpt020.

93. Schmull, M. & Thomas, F. M. Morphological and physiological reactions of young deciduous trees (Quercus robur L., Q. petraea [Matt.] Liebl., Fagus sylvatica L.) to waterlogging. Plant and Soil, 225(2000), 227–242 (2000). 10.1023/A:1026516027096.

94. Schultz, E. L., Hülsmann, L., Pillet, M. D., Hartig, F., Breshears, D. D., Record, S., Shaw, J. D., DeRose, R. J., Zuidema, P. A., & Evans, M. E. K. Climate-driven, but dynamic and complex? A reconciliation of competing hypotheses for species’ distributions. Ecology Letters, 25(1), 38–51 (2022). 10.1111/ele.13902.

95. Schurr, F. M., Pagel, J., Cabral, J. S., Groeneveld, J., Bykova, O., O’Hara, R. B., Hartig, F., Kissling, W. D., Linder, H. P., Midgley, G. F., Schröder, B., Singer, A., & Zimmermann, N. E. How to understand species’ niches and range dynamics: A demographic research agenda for biogeography. Journal of Biogeography, 39(12), 2146–2162 (2012). 10.1111/j.1365-2699.2012.02737.x.

96. Schwinning, S. Mechanisms determining the degree of size asymmetry in competition among plants. Oecologia, 113, 447–455 (1998). 10.1007/s004420050397.

97. Sharma, S., Andrus, R., Bergeron, Y., Bogdziewicz, M., Bragg, D. C., Brockway, D., Cleavitt, N. L., Courbaud, B., Das, A. J., Dietze, M., Fahey, T. J., Franklin, J. F., Gilbert, G. S., Greenberg, C. H., Guo, Q., Hille Ris Lambers, J., Ibanez, I., Johnstone, J. F., Kilner, C. L., Knops, J. M. H., et al. North American tree migration paced by climate in the West, lagging in the East. Proceedings of the National Academy of Sciences, 119(3), e2116691118 (2022). 10.1073/pnas.2116691118.

98. Stan Development Team. Stan Modeling Language Users Guide and Reference Manual. https://mc-stan.org (2022).

99. Stillhard, J., Hobi, M. L., Brang, P., Brändli, U.-B., Korol, M., Pokynchereda, V., & Abegg, M. Structural changes in a primeval beech forest at the landscape scale. Forest Ecology and Management, 504, 119836 (2022). 10.1016/j.foreco.2021.119836.

100. Sutherland, W. J., Freckleton, R. P., Godfray, H. C. J., Beissinger, S. R., Benton, T., Cameron, D. D., Carmel, Y., Coomes, D. A., Coulson, T., Emmerson, M. C., Hails, R. S., Hays, G. C., Hodgson, D. J., Hutchings, M. J., Johnson, D., Jones, J. P. G., Keeling, M. J., Kokko, H., Kunin, W. E., Lambin, X., et al. Identification of 100 fundamental ecological questions. Journal of Ecology, 101(1), 58–67 (2013). 10.1111/1365-2745.12025.

101. Ter Braak, C. J. & Prentice, I. C. A Theory of Gradient Analysis. In *Advances in Ecological Research*, volume 18, pp. 271–317. Elsevier (1988). 10.1016/S0065-2504(08)60183-X.

102. Thuiller, W., Münkemüller, T., Lavergne, S., Mouillot, D., Mouquet, N., Schiffers, K., & Gravel, D. A road map for integrating eco-evolutionary processes into biodiversity models. Ecology Letters, 16, 94–105 (2013). 10.1111/ele.12104.

103. Thuiller, W., Münkemüller, T., Schiffers, K. H., Georges, D., Dullinger, S., Eckhart, V. M., Edwards, T. C., Gravel, D., Kunstler, G., Merow, C., Moore, K., Piedallu, C., Vissault, S., Zimmermann, N. E., Zurell, D., & Schurr, F. M. Does probability of occurrence relate to population dynamics? Ecography, 37(12), 1155–1166 (2014). 10.1111/ecog.00836.

104. Tilman, D. Resource Competition and Community Structure. Number 17 in Monographs in Population Biology. Princeton University Press, Princeton, N.J. (1982).

105. Tomasella, M., Nardini, A., Hesse, B. D., Machlet, A., Matyssek, R., & Häberle, K.-H. Close to the edge: Effects of repeated severe drought on stem hydraulics and non-structural carbohydrates in European beech saplings. Tree Physiology, 39, 717–728 (2019). 10.1093/treephys/tpy142.

106. Tomppo, E., Gschwantner, T., & for Common, M. L. P. National Forest Inventories. Springer, Heidelberg Dordrecht London New York (2010).

107. Valladares, F. & Niinemets, Ü. Shade Tolerance, a Key Plant Feature of Complex Nature and Consequences. Annual Review of Ecology, Evolution, and Systematics, 39(1), 237–257 (2008). 10.1146/annurev.ecolsys.39.110707.173506.

108. Vehtari, A., Gelman, A., Simpson, D., Carpenter, B., & Bürkner, P.-C. Rank-Normalization, Folding, and Localization: An Improved R for Assessing Convergence of MCMC (with Discussion). Bayesian Analysis, 16(2), 667–718 (2021). 10.1214/20-BA1221.

109. Wagner, S., Collet, C., Madsen, P., Nakashizuka, T., Nyland, R. D., & Sagheb-Talebi, K. Beech regeneration research: From ecological to silvicultural aspects. Forest Ecology and Management, 259(11), 2172–2182 (2010). 10.1016/j.foreco.2010.02.029.

110. Watkinson, A. Density-dependence in single-species populations of plants. Journal of Theoretical Biology, 83(2), 345–357 (1980). 10.1016/0022-5193(80)90297-0.

111. Watkinson, A. R. On the Abundance of Plants Along an Environmental Gradient. The Journalof Ecology, 73(2), 569 (1985). 10.2307/2260494.

112. Whittaker, R. H. Gradient Analysis of Vegetation. Biological Reviews, 42(2), 207–264 (1967). 10.1111/j.1469-185X.1967.tb01419.x.

113. Wilkens, J. F. & Wagner, S. Empirical survival model for European beech (Fagus sylvatica L.) seedlings in response to interactive resource gradients and (a-) biotic conditions within an experimental canopy gap study. Forest Ecology and Management, 499, 119627 (2021). 10.1016/j.foreco.2021.119627.

114. Wood, S. Mgcv: Mixed GAM Computation Vehicle with Automatic Smoothness Estimation. https://CRAN.R-project.org/package=mgcv (2021).

115. Young, T. P., Petersen, D. A., & Clary, J. J. The ecology of restoration: Historical links, emerging issues and unexplored realms. Ecology Letters, 8(6), 662–673 (2005). 10.1111/j.1461-0248.2005.00764.x.

116. Zhu, K., Woodall, C. W., Ghosh, S., Gelfand, A. E., & Clark, J. S. Dual impacts of climate change: Forest migration and turnover through life history. Global Change Biology, 20(1), 251–264 (2014). 10.1111/gcb.12382.

117. Zurell, D., Grimm, V., Rossmanith, E., Zbinden, N., Zimmermann, N. E., & Schröder, B. Uncertainty in predictions of range dynamics: Black grouse climbing the Swiss Alps. Ecography, 35(7), 590–603 (2012). 10.1111/j.1600-0587.2011.07200.x.

118. Zurell, D., Thuiller, W., Pagel, J., Cabral, J. S., Münkemüller, T., Gravel, D., Dullinger, S., Normand, S., Schiffers, K. H., Moore, K. A., & Zimmermann, N. E. Benchmarking novel approaches for modelling species range dynamics. Global Change Biology, 22(8), 2651–2664 (2016). 10.1111/gcb.13251.

