## Supplementary Material for "How do sapling and adult demography explain beech predominance along environmental gradients in Central European forests?"


- **Lukas Heiland (corresponding author)**, University of Bayreuth, Bayreuth Center of Ecology and Environmental Research (BayCEER), Ecosystem Analysis and Simulation (EASI) Lab, Dr.-Hans-Frisch-Str. 1–3, 95448 Bayreuth, Germany. Theoretical Ecology, Universität Regensburg, Universitätsstraße 31, 93053 Regensburg, Germany.. ORCiD: https://orcid.org/0000-0002-0570-2138.
- **Lisa Hülsmann**, University of Bayreuth, Bayreuth Center of Ecology and Environmental Research (BayCEER), Ecosystem Analysis and Simulation (EASI) Lab, Dr.-Hans-Frisch-Str. 1–3, 95448 Bayreuth, Germany.. ORCiD: https://orcid.org/0000-0003-4252-2715.

### A Supplementary Methods

#### A.1 JAB model description

*Adapted from* Heiland et al. (2023).

The JAB model is a dynamic model with competition effects that includes an explicit juvenile stage (Figure S1; for code see `https://zenodo.org/records/8032461`). The JAB model describes populations of species that are logistically limited by the same resource (Gause, 1932) but differs from a classical competitive Lotka-Volterra model in four main aspects: (1) size-structured populations, (2) basal area growth, (3) reduced complexity of competition effects, (4) influx from outside the populations.

In the JAB model, populations are structured into three size stages that interact: the juvenile stage J, representing the understory, and the stages A and B, jointly representing the overstory (Figure S1). Partitioning tree populations into understory and overstory, we can express asymmetric competition between the two fundamentally different forest layers (Schwinning, 1998): the understory is affected by the shading of the overstory, while the overstory is directly exposed to light and unaffected by the understory (Valladares & Niinemets, 2008; Angelini et al., 2015; Cordonnier et al., 2019; De Lombaerde et al., 2019). The overstory (BA) is divided into A and B to enable conversion between measures of tree abundance in the sapling stage J, which is quantified as a count density, and the final stage B, which is represented in terms of basal area. The count density in J reflects the common measure of sapling inventories in NFI (see 2.2), whereas the basal area in the overstory stage B is a common measure for timber growth and competition (Biging & Dobbertin, 1992). The size stage A functions as an intermediary between J (only counts) and B (only basal area) by having counts that are converted to basal area with a conversion factor. The JAB model represents growth as transition rates in absence of competition: from the understory stage J to the intermediary stage A (parameter $g$) and from A to B (parameter $h$). In addition to transitions from A, the final stage B has intrinsic basal area growth (parameter $b$; Table 1).

To reduce the complexity of competition compared to a full Lotka-Volterra model, we represented only the differences in species' response to competition, assuming a similar competition effect among species (simplifying from a matrix of $n^2$ parameters to parameter vectors of $n$ competing species). More specifically, species are affected by the competition from the sum of the basal area of all species within their respective layers, i.e. they have a different competitive response to the sum of all inter- and intraspecific competition (vectors of species-specific parameters $c_J$, $c_A$, $c_B$). In applying this competition structure, we assume that the difference in competitive response between species is much more important than their difference in competitive effect (Tilman, 1982; Goldberg & Landa, 1991; Goldberg, 1996). The asymmetric competition from the overstory BA on J is represented by the "shading" parameter $s$.

In the JAB model, there are two different sources of seedling recruitment: (1) local recruitment that is proportional to the local conspecific basal area (parameter $r$), and (2) external seedling input. The external seedling input $L_p$ represents all long-term persistence of diaspores and long-distance dispersal into a subpopulation that is not not explained by the local conspecific basal area of a plot $p$. It is proportional to a measure of long-term and large-scale distribution of the species $\mathfrak{B}_p$ with the coefficient and parameter $l$ (detailed in A.2).

Based on these principles, we implemented the JAB model as a discrete-time iteration rule in stan, with hyperbolic density dependence (see Watkinson, 1980; Ellner, 1984; Levine & Rees, 2004), similar to a Lotka Volterra-type model formulation in Din (2013). The iteration rule comprises a set of four equations (Equations S1–S4) that relate states at year $t+1$ to states at year $t$. Here, in accordance with the software implementation, we provide a vectorized formulation of the model, where all variables, including the parameters, and the stages $J$ [$\mathrm{ha}^{-1}$], $A$ [$\mathrm{m}^2\ \mathrm{ha}^{-1}$], $B$ [$\mathrm{m}^2\ \mathrm{ha}^{-1}$], and $BA$ [$\mathrm{m}^2\ \mathrm{ha}^{-1}$] are vectors with length $n$ (number of species). These vectors are operated on with element-wise multiplication $\odot$ and division $\oslash$; the operator $\mathrm{sum}$ reduces the stages to a scalar, representing the total abundance of a stage across species.

$$J_{t+1} = L_p + r \odot BA_t + (J_t - g \odot J_t) \oslash [1 + c_J\ \mathrm{sum}(J_t) + s\ \mathrm{sum}(BA_t)] \tag{S1}$$

$$A_{t+1} = g \odot J_t \oslash [1 + c_J\ \mathrm{sum}(J_t) + s\ \mathrm{sum}(BA_t)] + (A_t - h \odot A_t) \oslash [1 + c_A\ \mathrm{sum}(BA_t)] \tag{S2}$$

$$B_{t+1} = h \odot A_t \cdot \beta_{uA} \oslash [1 + c_A\ \mathrm{sum}(BA_t)] + (1 + b) \odot B_t \oslash [1 + c_B\ \mathrm{sum}(BA_t)] \tag{S3}$$

$$BA_{t+1} = A_{t+1} \odot \beta_{mA} + B_{t+1} \tag{S4}$$

All parameters ($r, c_J, s, g, c_A, h, b, c_B$, and $L_p = l\mathfrak{B}_p$) are generally assumed to be positive, so that all model states are strictly positive at any time (in fitting the model, this will be ensured by exponentiating the parameters sampled on a log-scale; Section 2.4.1). The four equations, representing size stages, are coupled through states of other stages, so that changes in one state propagate in discrete time steps, e.g., from BA to J to A to B. The fractions of trees that survive and grow from J to A and from A to B are expressed by the transition rates $g$ and $h$ ($\in (0,1)$), respectively. The counts in A are transformed to basal area at two different occasions: (1) In the transition to B, the counts are converted by factoring in the basal area of one tree at the threshold between A and B, a species-independent scalar factor $\beta_{uA}$, which is dependent on the threshold diameter at breast height (dbh; Equation S3); (2) the combined basal area BA is calculated by multiplying the counts within A with a vector of the corresponding mean basal areas $\beta_{mA}$, which are species-specific constants from the data (Equation S4; *Fagus*: $0.015687\mathrm{m}^2$; *others* $0.016069\mathrm{m}^2$).

All stages (J, A, and B) are logistically limited by interspecific tree density: The sapling stage J is limited by the competitive effect from the total basal area across species BA ($s$) and from the total counts within the same stage ($c_J$), while the stages A and B are only limited by the competitive effect of BA ($c_A$, $c_B$). This limitation is implemented by dividing the states with a denominator that is slightly greater than 1: $[1 + c_J\ \mathrm{sum}(J_t) + s\ \mathrm{sum}(BA_t)]$ and $[1 + c_{A,B}\ \mathrm{sum}(BA_t)]$, for understory and overstory respectively. The fractions of trees that transitions to the next stage, i.e. $g \cdot J$ and $h \cdot A$, are also limited by tree density so that they cannot exceed the number of trees in the current stage that is affected by limitation, e.g., the trees that transition from A to B $h \odot A_t \oslash [1 + c_A\ \mathrm{sum}(BA_t)]$ cannot exceed $A_t \oslash [1 + c_A\ \mathrm{sum}(BA_t)]$. Similarly in stage B, the density-dependent increment $B_t \odot b \oslash [1 + c_B\ \mathrm{sum}(BA_t)]$ cannot exceed the state-conserving term $B_t \oslash [1 + c_B\ \mathrm{sum}(BA_t)]$ (with $b \leq 1$). Not making $B_t \odot b$ density-dependent would allow constellations where $b \sim 1$ and $c_B >> 0$ so that most of the state conservation would come from $B_t \odot b$. As a result, the competitive parameters $c_J$, $s$, $c_A$, and $c_B$ describe, how species are differently affected by density of the respective forest layers ($\mathrm{sum}\ J_t$ and $\mathrm{sum}\ BA_t$) and the parameters $g$, $h$ and $b$ describe growth and survival in the absence of competition.

Overall, this leads to a system of populations that are in an arms race from a disturbed state towards a competitive equilibrium: Depending on their seedling recruitment ($L_p$ and $r$), through transition ($g$, $h$) and net basal area increment ($b$) populations can intrinsically only grow or stagnate. This assumes that density-independent mortality is negligible in J and A, and that density-independent mortality in B is included in $b$ and does not exceed basal area growth in the long term. Populations can, however, decline through interspecific competition ($c_J$, $s$, $c_A$, $c_B$). These properties enable the JAB model to extrapolate species composition under the assumption that species composition is mainly determined by a competitive equilibrium (Ellenberg, 1963).

#### A.2 Species' regional abundance for predicting seedling input

*Reproduced from* Heiland et al. (2023).

To derive the regional species abundance $\mathfrak{B}_p$ used to estimate the external seedling input (parameter $l$, see Section A.1) we used thin plate spline regressions with the geographic coordinates as predictors to interpolate the species-specific basal area per hectare on the German NFI grid (see Appendix of Heiland et al., 2023 for estimate table and plots).

For fitting the thin plate splines from the regular standard grid of the German NFI (4 km; see 2.2), which is homogeneous across Federal states, we selected only the basal area records from the last survey (2012), because the grid had changed over surveys. We completed non-forested clusters of the regular grid, and also all of the four plots within clusters, with zero observations to obtain an unbiased sample of the geographical density of tree species. To obtain a response variable for each set of coordinates, we calculated the species abundances per cluster by averaging the basal area per hectare above the sampling threshold (BA) over the four plots. This average basal area per hectare on the completed grid was spatially smoothed with a two-dimensional geographic thin plate spline. Based on the coordinates, a "spline on sphere" was fitted with the `gam()` function from the package mgcv (`k = 200`; version 1.8-39; Wood, 2021), where the average basal area per hectare was the response. The response variable had been rounded to an integer to be able to fit a negative binomial response. The predicted plot-specific regional species abundances $\mathfrak{B}_p$ were used as a proxy for the probability of seedling input, which factors into the JAB model with a species-specific effect size $l$, $L_p = \mathfrak{B}_p\, l$. By using a strictly positive negative binomial response for $\mathfrak{B}_p$, it was made sure that plots were not excluded from invasion in the JAB model.

#### A.3 Offsets for scaling area-standardized model states to observed counts

*Adapted from* Heiland et al. (2023).

To model tree abundances from varying sampling areas with a count process in the likelihood of the JAB model (Section 2.4.3), we used different offsets. In general, the offsets $o$ convert the model states $\hat{x}$, which are abundances standardized per hectar, to the scale of observed counts $x$:

$$x \sim o\,\hat{x} \tag{S5}$$

Different offsets were used depending on the type of the abundances in the model and the corresponding data: for the special offsets that scaled the basal area states in the model to count data from the angle count method see A.3.1; for the offsets that scaled counts in the model to counts on different fixed areas see A.3.2. Because the sampling protocol of the German NFI made observation areas dependent on size, the offsets for zero observations had to be derived separately (see A.3.3).

These offsets account for the varying sampling intensity that also affects the variance of observations. Furthermore, using a count process with offsets is (1) preferable over modelling a continuous response for the basal area because it reflects the actual observation process with discrete numbers of trees (even when being multiplied with a tree-specific basal area), and (2) preferable over upscaling the small sampling areas to a common area because this would break distributional assumptions by deflating small counts in the data, e.g. when a plot size is one fourth of the common standard area 1 hectare, the smallest measured count per hectare would be 4.

##### A.3.1 Offsets for the angle count method

Count data from the angle count method (size stage A in the German NFI) were related to the count states in the JAB model by scaling with area offsets. In angle count sampling, each tree has a specific sampling area $a$ dependent on its dbh with $a = \pi\, k^2\, dbh^2$, where $k$ is a constant for the sampling angle (here, $k = 25$). Thus, for the sampling area of a plot, which corresponds to the actually observed trees on a plot we use the weighted mean $a_p = \sum_{i=1}^{n} w_i a_i \,/\, \sum_{i=1}^{n} w_i$, where the weights $w_i$ are the respective area-standardized counts per hectare. This weighted mean has the property that—together with the actually observed counts on a plot $c_p$—it conveys equivalent information as the sum of all area-standardized counts per hectare on a plot $c_A = \frac{c_p}{a_p}$ [ha$^{-1}$]. The area $a_p$ was used as the offset for size stage A ($o$ in Equation S5).

To relate count data from the angle count method to the basal area state in the model (size stage B in the German NFI), we used an offset factor that not only includes the sampling area $a_p$ but transforms the basal area per hectare in the JAB model to counts. This was achieved with an offset $o$ that also included the total observed basal area $ba_p$ on a plot: $o = a_p \, \frac{c_p}{ba_p}$ [ha m$^{-2}$]. By multiplying the model state $\hat{x}$ [m$^2$ ha$^{-1}$] with this offset we transformed the model state to counts, by including the data on basal area ("how many counts are one unit of basal area?") on the right hand side of the model statement: $x \sim \; a_p \frac{c_p}{ba_p} \, \hat{x}$.

##### A.3.2 Offsets for sapling counts on fixed-area plots

Sapling counts on fixed-area plots (size class J) were also standardized using area offsets. In the German NFI data, size class J consists of multiple size classes with different sampling areas, which also change between surveys (Table S2). As in the angle count data, we calculated a weighted mean area per plot, where the different sampling areas were weighted by the counts per area within the corresponding size classes. The weighted mean area per NFI plot was used as an offset to relate the sapling counts to the area-standardized counts of J in the JAB model [ha$^{-1}$].

##### A.3.3 Offsets for zero observations

Because the sampling protocol of the German NFI made observation areas dependent on size, the observation area was not readily available when there were no observations for a stage on a plot. For zero observations in the angle count data in the German NFI (size classes A and B), we used the sampling area of the other species, or, if both species had zero observations, the mean sampling area per size class and survey as an offset (for a test of the robustness of this approach, see below). To be able to compute a mean area for the largest size class B, we truncated the angle count data to a maximum sampling radius of 15 m. The truncation included dropping all trees outside the sampling radius, which is about the 98th percentile of tree counts in A and B and assigning the new maximum sampling area to all trees that were within the maximum radius but had a dbh that was originally corresponding to a sampling area greater than the new maximum area (dbh $>$ 60 cm). Truncating the angle count data to a fixed maximum area removes a source of bias (missed tree observations at higher radii) at the cost of moderate added variance (Berger et al., 2020), and furthermore leads to a finite sampling area that can be averaged over. The mean sampling area per size class (A and B) was calculated by dividing the definite integral from 0 up to the respective maximum radius—by the maximum radius. This mean area was used as an offset for zero observations in size class A. To obtain an offset for zero observations in size class B, we also needed an additional factor that scaled the basal area to the scale of observed counts. Instead of multiplying the area with the plot-specific factor $\frac{c_p}{ba_p}$ (as for non-zero observations, see A.3.1), we included the mean of the factor per survey and additionally per species to differentiate between their average basal areas.

Zero observations in the sapling counts of the size class J in the German NFI were assigned an equally weighted mean of all possible sampling areas for the respective set of smaller size classes within the corresponding survey (Table S2).

### B Supplementary Results

#### B.1 Estimates of demographic rates

The fitted paraboloid parameters ($\epsilon$, $\kappa$, $\zeta$; Section 2.4.2) determined the JAB model parameter values at the subpopulation level ($r$, $c_J$, $s$, $g$, $c_A$, $h$, $b$, and $c_B$). The resulting variation of subpopulation-level rates across environments (*environmental variation*; Section 2.4.2), was primarily influenced by the extreme values $\epsilon$ (Table 2).

Notably, the extreme values and means of the environmental variation showed the most differentiation between *Fagus* and *others* at the sapling stage J compared to the later stages (Table 2). Specifically, *others'* saplings exhibited competition response that was orders of magnitude higher than *Fagus'*, both in terms of competition from the overstory ($s$) and within-sapling competition ($c_J$). while *Fagus* had much lower estimates for seedling recruitment ($r$) and external seedling input ($L_p$).

In the overstory stages A and B, *others* still had greater response to competition, but the species differences decreased as tree size increased from A ($c_A$) to B ($c_B$).

Regarding growth and transition rates in the absence of competition ($g$, $h$, $b$), the species differences varied across size classes. *Fagus* exhibited a lower fraction of saplings transitioning to stage A ($g$) but a higher fraction transitioning from stage A to stage B ($h$). The net basal area increment ($b$) was smaller for *Fagus* compared to *others*. These species relations generally held true for density-dependent growth terms (Section 2.1), except for the transition term $\frac{g}{1+s\,\mathrm{sum}(BA)+c_J\,\mathrm{sum}(J)}$, which was greater for *Fagus* but only at the equilibrium.

### References for Supplementary Methods

Angelini, A., Corona, P., Chianucci, F., & Portoghesi, L. Structural attributes of stand overstory and light under the canopy. *Annals of Silvicultural Research*, **39**(1), 23–31 (2015). http://dx.doi.org/10.12899/asr-993.

Berger, A., Gschwantner, T., & Schadauer, K. The effects of truncating the angle count sampling method on the Austrian National Forest Inventory. *Annals of Forest Science*, **77**(1), 16 (2020). http://dx.doi.org/10.1007/s13595-019-0907-y.

Biging, G. S. & Dobbertin, M. A comparison of distance-dependent competition measures for height and basal area growth of individual conifer trees. *Forest Science*, **38**(3), 695–720 (1992). http://dx.doi.org/10.1093/forestscience/38.3.695.

Cordonnier, T., Smadi, C., Kunstler, G., & Courbaud, B. Asymmetric competition, ontogenetic growth and size inequality drive the difference in productivity between two-strata and one-stratum forest stands. *Theoretical Population Biology*, **130**, 83–93 (2019). http://dx.doi.org/10.1016/j.tpb.2019.07.001.

De Lombaerde, E., Verheyen, K., Van Calster, H., & Baeten, L. Tree regeneration responds more to shade casting by the overstorey and competition in the understorey than to abundance per se. *Forest Ecology and Management*, **450**, 117492 (2019). http://dx.doi.org/10.1016/j.foreco.2019.117492.

Din, Q. Dynamics of a discrete Lotka-Volterra model. *Advances in Difference Equations*, **2013**(95), 1–13 (2013). http://dx.doi.org/10.1186/1687-1847-2013-95.

Ellenberg, H. *Vegetation Mitteleuropas Mit Den Alpen. In Kausaler, Dynamischer Und Historischer Sicht.*, volume IV/2 of *Einführung in Die Phytologie.* Ulmer, Stuttgart, 1 edition (1963).

Ellner, S. Asymptotic behavior of some stochastic difference equation population models. *Journal Of Mathematical Biology*, **19**(2), 169–200 (1984). http://dx.doi.org/10.1007/BF00277745.

Gause, G. F. Experimental studies on the struggle for existence: I. Mixed population of two species of yeast. *Journal of experimental biology*, **9**(4), 389–402 (1932). http://dx.doi.org/10.1242/jeb.9.4.389.

Goldberg, D. E. Competitive ability: Definitions, contingency and correlated traits. *Philosophical Transactions of the Royal Society of London. Series B: Biological Sciences*, **351**(1345), 1377–1385 (1996). http://dx.doi.org/10.1098/rstb.1996.0121.

Goldberg, D. E. & Landa, K. Competitive Effect and Response: Hierarchies and Correlated Traits in the Early Stages of Competition. *The Journal of Ecology*, **79**(4), 1013 (1991). http://dx.doi.org/10.2307/2261095.

Heiland, L., Kunstler, G., Šebeň, V., & Hülsmann, L. Which demographic processes control competitive equilibria? Bayesian calibration of a size-structured forest population model. *Ecology and Evolution*, **13**(7), e10232 (2023). http://dx.doi.org/10.1002/ece3.10232.

Levine, J. M. & Rees, M. Effects of Temporal Variability on Rare Plant Persistence in Annual Systems. *The American Naturalist*, **164**(3), 350–363 (2004). http://dx.doi.org/10.1086/422859.

Schwinning, S. Mechanisms determining the degree of size asymmetry in competition among plants. *Oecologia*, **113**, 447–455 (1998). http://dx.doi.org/10.1007/s004420050397.

Tilman, D. *Resource Competition and Community Structure.* Number 17 in Monographs in Population

Biology. Princeton University Press, Princeton, N.J. (1982).

Valladares, F. & Niinemets, Ü. Shade Tolerance, a Key Plant Feature of Complex Nature and Consequences. *Annual Review of Ecology, Evolution, and Systematics,* **39**(1), 237–257 (2008). http://dx.doi.org/10.1146/annurev.ecolsys.39.110707.173506.

Watkinson, A. Density-dependence in single-species populations of plants. *Journal of Theoretical Biology,* **83**(2), 345–357 (1980). http://dx.doi.org/10.1016/0022-5193(80)90297-0.

Wood, S. Mgcv: Mixed GAM Computation Vehicle with Automatic Smoothness Estimation. https://CRAN.R-project.org/package=mgcv (2021).

### C Supplementary Tables

**Table S1:** Scope of the German NFI and plot selection.

| German NFI | |
|---|---|
| No. of surveys | 3 |
| ... years | 1986–1989/2000–2003/2011–2013 |
| No. of plots | 61666 |
| ... after selection | 795 |
| No. of clusters | 21574 |
| ... after selection | 795 |

**Table S2:** Size classes for saplings counts and corresponding radii of sampling circles in the three surveys of the German NFI (main years 1987, 2002, 2012). The size classes up to dbh 7 cm, which were consistently sampled across all three surveys, were assigned to the size class J in fitting the JAB model.

| | 1987 | 2002 | 2012 |
|---|---|---|---|
| height 20–50 cm | 1 m | 1 m | 1 m |
| height 50–130 cm | 2 m | 1.75 m | 2 m |
| height 130–$\infty$ cm and dbh 0–5 cm | 2 m | 1.75 m | 2 m |
| dbh 5–6 cm | 4 m | 1.75 m | 2 m |
| dbh 6–7 cm | 4 m | 1.75 m | 2 m |
| dbh 7–8 cm | 4 m | . | . |
| dbh 8–9 cm | 4 m | . | . |
| dbh 9–10 cm | 4 m | . | . |

**Table S3:** Soil water levels from Benning et al. (2016), ranking and conversion into pseudo-ratio scale. Translation according to Leuschner & Ellenberg (2017a).

| Category | Description | Pseudo-ratio value | Ellenberg levels | Translation |
|---|---|---|---|---|
| T1 | trocken | -6 | trocken | dry |
| T1-T2 | trocken bis mäßig trocken | -5 | | |
| T2 | mäßig trocken | -4 | mäßig trocken | moderately dry |
| T2-T3 | mäßig trocken bis mäßig frisch | -3 | | |
| T2-T4 | mäßig trocken bis frisch | -2.5 | | |
| T3 | mäßig frisch | -2 | mäßig frisch | moderately damp |
| T3-T4 | mäßig frisch bis frisch | -1 | | |
| S | sehr tief sitzende Staunässe | -0.5 | | |
| T3-S1 | mäßig frisch bis sehr schwach staunass / wechselfeucht | -0.25 | | |
| T4 | frisch | 0 | frisch | damp |
| S0 | tief sitzende Staunässe | 0.25 | | |
| T4-T5 | frisch bis sehr frisch | 0.5 | | |
| T4-S1 | frisch bis sehr schwach staunass / wechselfeucht | 0.7 | | |
| T5 | sehr frisch | 0.8 | | |
| G0 | grundfrisch | 1 | | |
| S1 | sehr schwach staunass / wechselfeucht | 1.25 | | |
| G1 | grundfrisch | 1.5 | | |
| S1-S2 | sehr schwach bis schwach staunass / wechselfeucht | 1.7 | | |
| S2 | schwach staunass / wechselfeucht | 1.8 | | |
| G2 | grundfeucht | 2 | (mâßig feucht) | (moderately moist) |
| S1-G2/3 | sehr schwach staunass / wechselfeucht bis grundfeucht | 2.2 | | |
| S1-S3 | sehr schwach bis mittel staunass / wechselfeucht | 2.3 | | |
| G2/3 | grundfeucht | 2.5 | | |
| S3 | mittel staunass / wechselfeucht | 2.7 | | |
| S3-G2/3 | mittel staunass / wechselfeucht bis grundfeucht | 2.8 | | |
| G3 | grundfeucht | 3 | | |
| S3-S4 | mittel bis stark staunass / wechselfeucht | 3.5 | | |
| G4 | feucht | 4 | feucht | moist |
| G4-G5 | feucht bis nass | 5 | | |
| S4 | stark staunass / wechselfeucht | 5.5 | | |
| G4-G6 | feucht bis nass | 5.8 | | |
| G5 | nass | 6 | nass | wet |
| S3-S6 | mittel bis äußerst staunass | 6.5 | | |
| S5 | sehr stark staunass / wechselnass | 7 | | |
| G6 | nass | 7.5 | | |
| S6 | äußerst staunass | 8 | sehr nass | very wet |

**Table S4:** Stratification of the two environmental gradients into seven intervals each (left: water level, top: pH) and the number of forest plots sampled within each of the 49 resulting bins.

| | (2.84,3.47] | (3.47,4.09] | (4.09,4.72] | (4.72,5.34] | (5.34,5.97] | (5.97,6.59] | (6.59,7.22] |
|---|---|---|---|---|---|---|---|
| (-6.01,-4.07] | 1 | 14 | 23 | 22 | 25 | 6 | 4 |
| (-4.07,-2.14] | 6 | 25 | 25 | 25 | 25 | 25 | 19 |
| (-2.14,-0.214] | 15 | 25 | 25 | 25 | 25 | 25 | 6 |
| (-0.214,1.71] | 15 | 25 | 25 | 25 | 25 | 25 | 6 |
| (1.71,3.64] | 4 | 13 | 25 | 25 | 25 | 12 | 3 |
| (3.64,5.57] | 3 | 21 | 25 | 25 | 25 | 21 | 2 |
| (5.57,7.51] | 0 | 4 | 6 | 6 | 7 | 5 | 1 |

**Table S5:** Taxa composition of the German NFI on the selected plots. All observed taxa are listed, ranked by the mean basal area per plot. In addition, the mean percentage of the total plot basal area is given.

| | Taxon | Mean basal area $m^2$ $ha^{-1}$ | Mean percentage of the total basal area |
|---|---|---|---|
| 1 | Picea abies | 7.852 | 28.972% |
| 2 | Pinus sylvestris | 5.714 | 18.566% |
| 3 | Fagus sylvatica | 3.325 | 13.102% |
| 4 | Quercus petraea | 1.761 | 6.754% |
| 5 | Quercus robur | 1.249 | 4.522% |
| 6 | Pseudotsuga menziesii | 0.886 | 3.391% |
| 7 | Betula pendula | 0.692 | 3.161% |
| 8 | Alnus glutinosa | 0.739 | 2.581% |
| 9 | Fraxinus excelsior | 0.562 | 2.478% |
| 10 | Larix decidua | 0.637 | 2.356% |
| 11 | Carpinus betulus | 0.555 | 1.778% |
| 12 | Quercus spp. | 0.561 | 1.717% |
| 13 | Acer pseudoplatanus | 0.235 | 1.318% |
| 14 | Abies alba | 0.281 | 1.048% |
| 15 | Prunus avium | 0.218 | 0.899% |
| 16 | Alnus spp. | 0.237 | 0.760% |
| 17 | Salix spp.. | 0.14 | 0.691% |
| 18 | Populus tremula | 0.187 | 0.661% |
| 19 | Tilia spp. | 0.181 | 0.649% |
| 20 | Quercus rubra | 0.137 | 0.589% |
| 21 | Populus nigra | 0.135 | 0.460% |
| 22 | Robinia pseudoacacia | 0.17 | 0.389% |
| 23 | Acer campestre | 0.09 | 0.382% |
| 24 | Populus | 0.147 | 0.366% |
| 25 | Larix kaempferi | 0.081 | 0.341% |
| 26 | Pinus nigra | 0.103 | 0.236% |
| 27 | Larix spp. | 0.075 | 0.218% |
| 28 | Acer platanoides | 0.03 | 0.195% |
| 29 | Alnus incana | 0.035 | 0.183% |
| 30 | Pinus strobus | 0.039 | 0.156% |
| 31 | Ulmus spp. | 0.034 | 0.124% |
| 32 | Populus alba | 0.073 | 0.093% |
| 33 | other coniferous (German NFI) | 0.029 | 0.088% |
| 34 | Sorbus aucuparia | 0.012 | 0.083% |
| 35 | Aesculus hippocastanum | 0.024 | 0.078% |
| 36 | other deciduous (German NFI) | 0.017 | 0.070% |
| 37 | Populus trichocarpa x maximoviczii | 0.03 | 0.065% |
| 38 | Sorbus aria | 0.012 | 0.060% |
| 39 | Picea sitchensis | 0.01 | 0.058% |
| 40 | Sorbus torminalis | 0.017 | 0.051% |
| 41 | Thuja spp. | 0.02 | 0.041% |
| 42 | Prunus padus | 0.007 | 0.038% |
| 43 | Betula pubescens | 0.005 | 0.035% |
| 44 | Ilex aquifolium | 0.01 | 0.032% |
| 45 | Frangula alnus | 0.005 | 0.030% |
| 46 | Castanea sativa | 0.005 | 0.027% |
| 47 | Pyrus communis | 0.007 | 0.027% |
| 48 | other Pinus (German NFI) | 0.008 | 0.026% |
| 49 | Sorbus spp. | 0.005 | 0.019% |
| 50 | Abies grandis | 0.003 | 0.018% |
| 51 | Juglans | 0.003 | 0.016% |
| 52 | Malus sylvestris | 0.002 | 0.004% |

**Table S6:** Posterior distributions of the dispersion parameter $\phi$ and bulk effective sample size (ESS). There are different levels of uncertainty per stage and survey.

| | Level | Species | Posterior | bulk ESS |
|---|---|---|---|---|
| $\phi$ | J Initial state | Fagus | 10170. ± 152700. | 3962.175 |
| | J Initial state | others | 21850. ± 173200. | 4114.269 |
| | J 2nd survey | Fagus | 0.3863 ± 0.05948 | 8763.802 |
| | J 2nd survey | others | 0.2342 ± 0.01473 | 7526.126 |
| | J 3rd survey | Fagus | 0.2050 ± 0.01912 | 9997.296 |
| | J 3rd survey | others | 0.2066 ± 0.01238 | 9309.075 |
| | A Initial state | Fagus | 5157. ± 20760. | 3520.311 |
| | A Initial state | others | 7000. ± 36340. | 4753.705 |
| | B Initial state | Fagus | 9725. ± 165700. | 3703.357 |
| | B Initial state | others | 12330. ± 65700. | 4243.675 |
| | A 2nd/3d survey | Fagus | 3.623 ± 1.010 | 6371.580 |
| | A 2nd/3d survey | others | 1.980 ± 0.1897 | 8463.309 |
| | B 2nd/3d survey | Fagus | 2.485 ± 0.3213 | 7960.757 |
| | B 2nd/3d survey | others | 3.964 ± 0.2868 | 6709.977 |

**Table S7:** Posterior distributions of initial and equilibrium abundances (stages J and A [count $ha^{-1}$]; B and total BA [$m^2$ $ha^{-1}$]) and species' predominance (mean $\pm$ standard deviation across subpopulations and HMC samples). Equilibrium states include counterfactual simulations, where the rate $s$ has been switched between *Fagus* and *others* or where the respective other species does not regenerate.

| | | Fagus | others |
|---|---|---|---|
| J | initial state data | $817.05 \pm 4911.0$ | $4414.5 \pm 11196.$ |
| | initial model state | $827.12 \pm 4912.4$ | $4436.1 \pm 11199.$ |
| | equilibrium state | $7789.7 \pm 4342.6$ | $3218.9 \pm 1575.8$ |
| A | initial state data | $43.744 \pm 187.64$ | $312.88 \pm 525.15$ |
| | initial model state | $45.227 \pm 187.23$ | $313.88 \pm 524.46$ |
| | equilibrium state | $550.96 \pm 295.18$ | $76.920 \pm 58.745$ |
| B | initial state data | $2.0691 \pm 5.9266$ | $14.140 \pm 15.510$ |
| | initial model state | $2.1133 \pm 6.0062$ | $14.258 \pm 15.678$ |
| | equilibrium state | $39.382 \pm 31.583$ | $7.7577 \pm 10.855$ |
| BA | initial state data | $2.6943 \pm 7.0598$ | $18.590 \pm 16.974$ |
| | initial model state | $2.8227 \pm 7.2233$ | $19.302 \pm 17.430$ |
| | equilibrium state | $48.025 \pm 34.253$ | $8.9937 \pm 11.752$ |
| | eq. sum across species | $57.019 \pm 28.099$ | . |
| | eq. with switched $s$ | $8.0109 \pm 20.301$ | $32.853 \pm 12.054$ |
| | eq. without regeneration of resp. other | $58.395 \pm 30.592$ | $38.786 \pm 2.8381$ |
| Fraction of basal area | initial model state | $0.20731 \pm 0.31625$ | $0.79269 \pm 0.31625$ |
| | equilibrium state | $0.78654 \pm 0.30819$ | $0.21346 \pm 0.30819$ |
| | eq. with switched $s$ | $0.16222 \pm 0.30936$ | $0.16222 \pm 0.30936$ |

### D Supplementary Figures

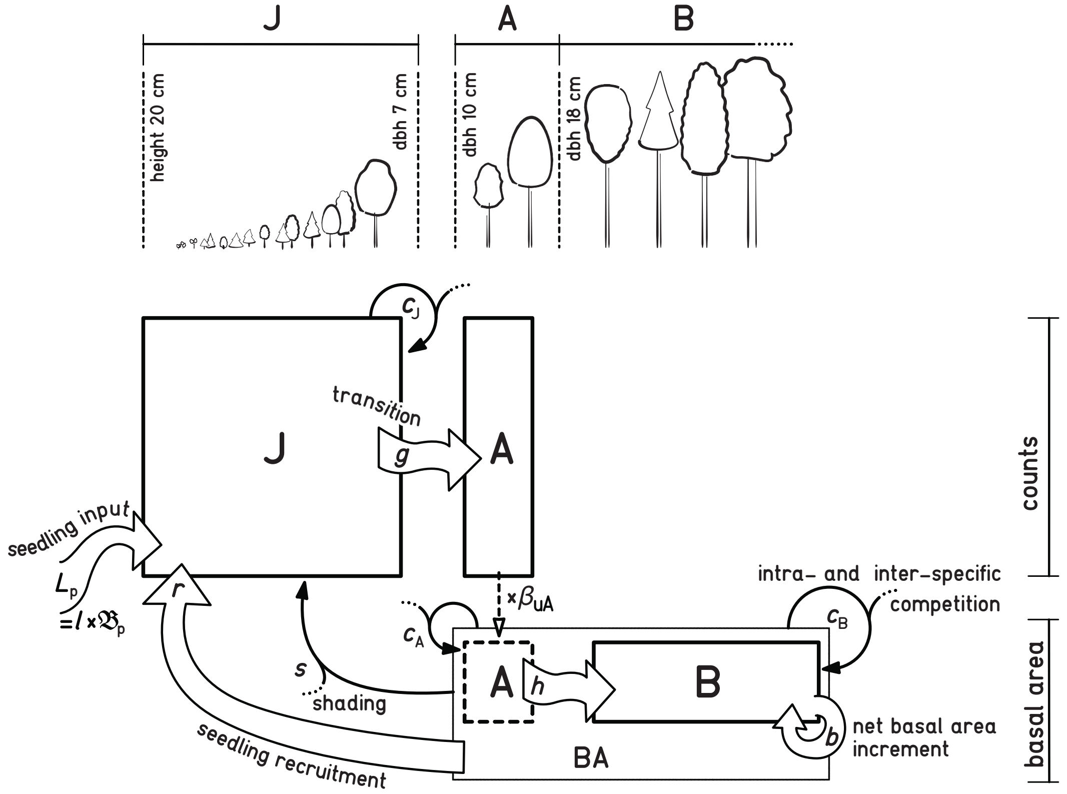


**Figure S1:** The JAB model represents three different size stages of a tree population (J, A, B) and demographic processes, including the competitive interactions between size stages and multiple species. There are processes that contribute to population growth: seedling input into the population from temporal or spatial dispersal $L_p$ (dependent on regional basal area $\mathfrak{B}_p$), within-population seedling regeneration $r$, transitions from a smaller to a larger stage ($g$, $h$), and the basal area growth $b$. The processes that limit population growth comprise the competition effect from the total sum of the sapling stage J on species-specific J ($c_J$), the competition from the sum of A and B, i.e. the total basal area of all species BA, on A and B ($c_A$, $c_B$), as well as the asymmetric competition from BA on J ("shading" effect $s$). The choice of thresholds between size stages is informed by the size classes in the data. In the German NFI small trees between height 20 cm and dbh 7 cm were counted, while trees with dbh $>$ 10 cm were additionally measured in terms of basal area, so that intermediary size stage A acts as a mediator beween count data for J and the basal area data for B. The factor $\beta_{uA}$, i.e. the upper basal area of a tree in A, converts A from counts to basal area.

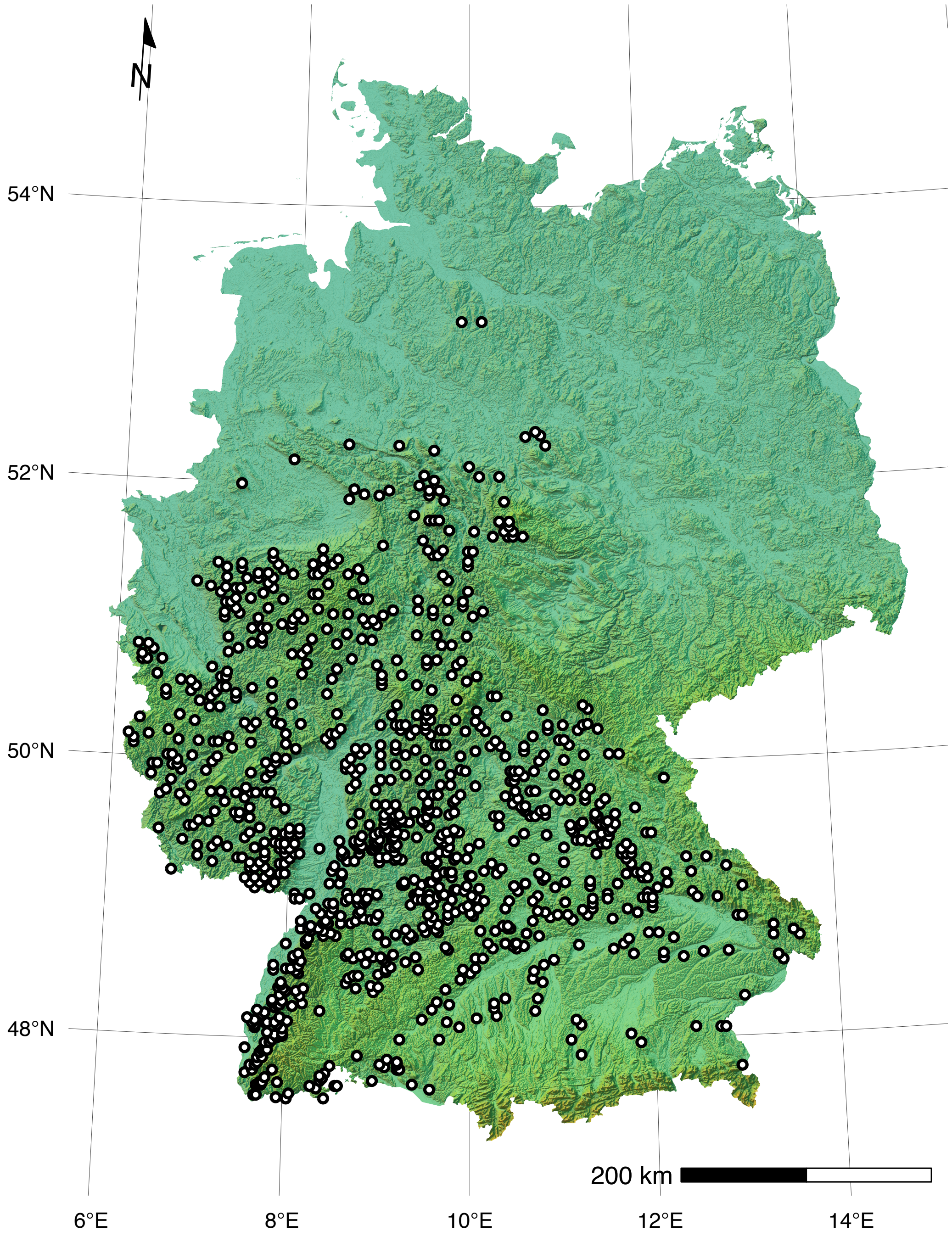


**Figure S2:** Locations of the 795 randomly selected NFI clusters in Germany. Each cluster consists of one to four sampling plots, depending on whether the location was forested. One random plot per cluster was selected if its elevation was at 100–600 m, if the observation period included all of the three surveys (1987, 2002, and 2012; which excludes former East Germany), and if there were no records of management within the period. Each selected plot (cluster) is represented by one subpopulation in the JAB model. The map projection is Lambert Conformal Conic, the color shading indicates elevation and relief.

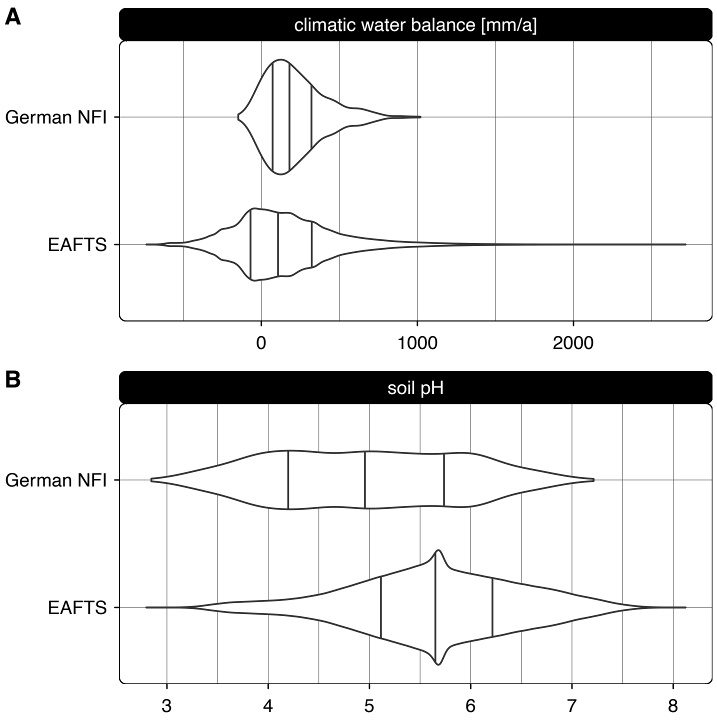


**Figure S3:** Environmental range of the German NFI and of *Fagus sylvatica* occurrence according to The European Atlas of Forest Tree Species (de Rigo et al., 2016). The German NFI has a narrower range with regard to both top soil pH in $CaCl_2$ (**A**) and climatic water balance (**B**). Lines correspond to quartiles.

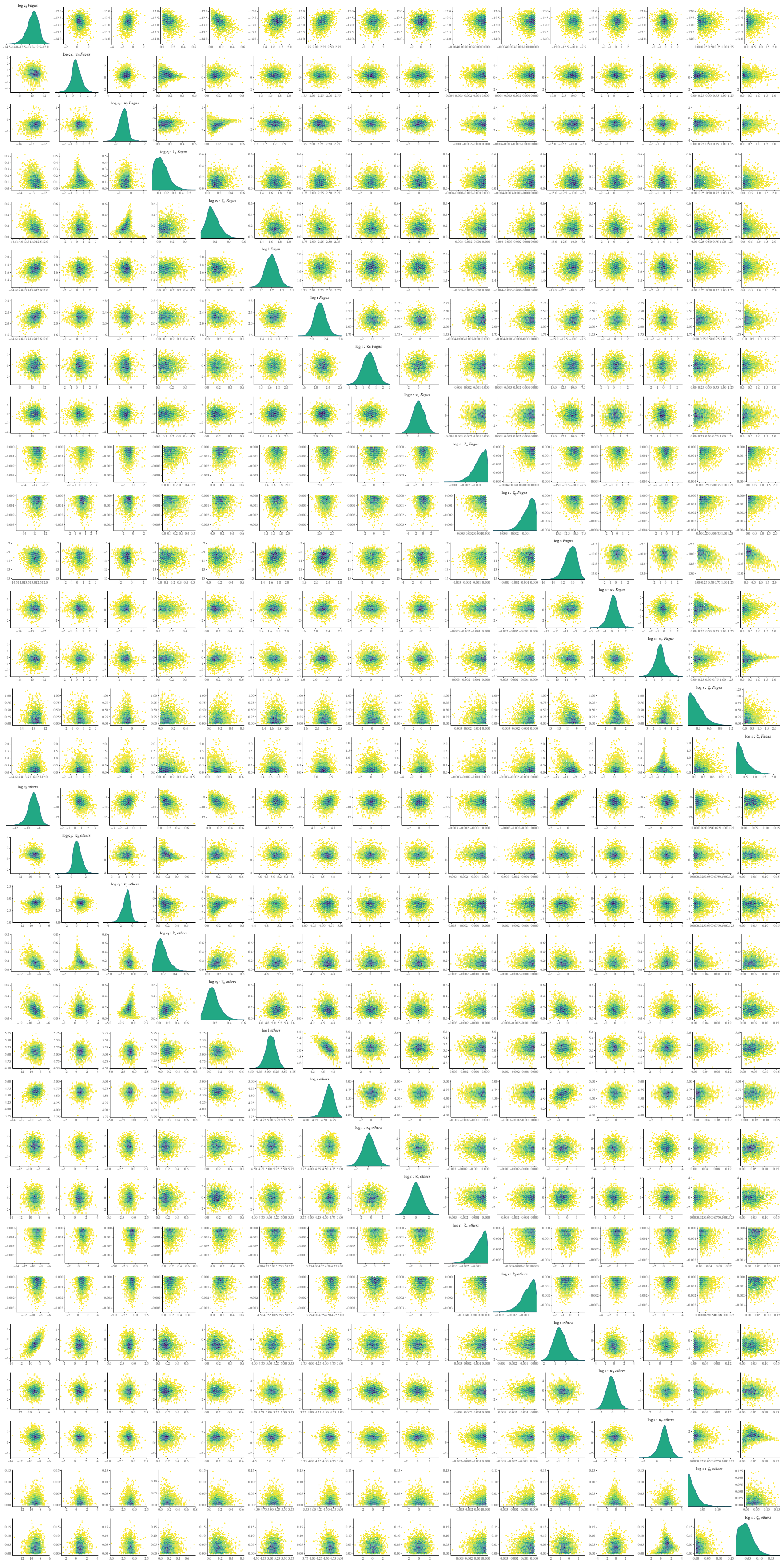

**Figure S4:** Pairs of paraboloid parameter correlations within stage J, separately for *Fagus* and *others*.

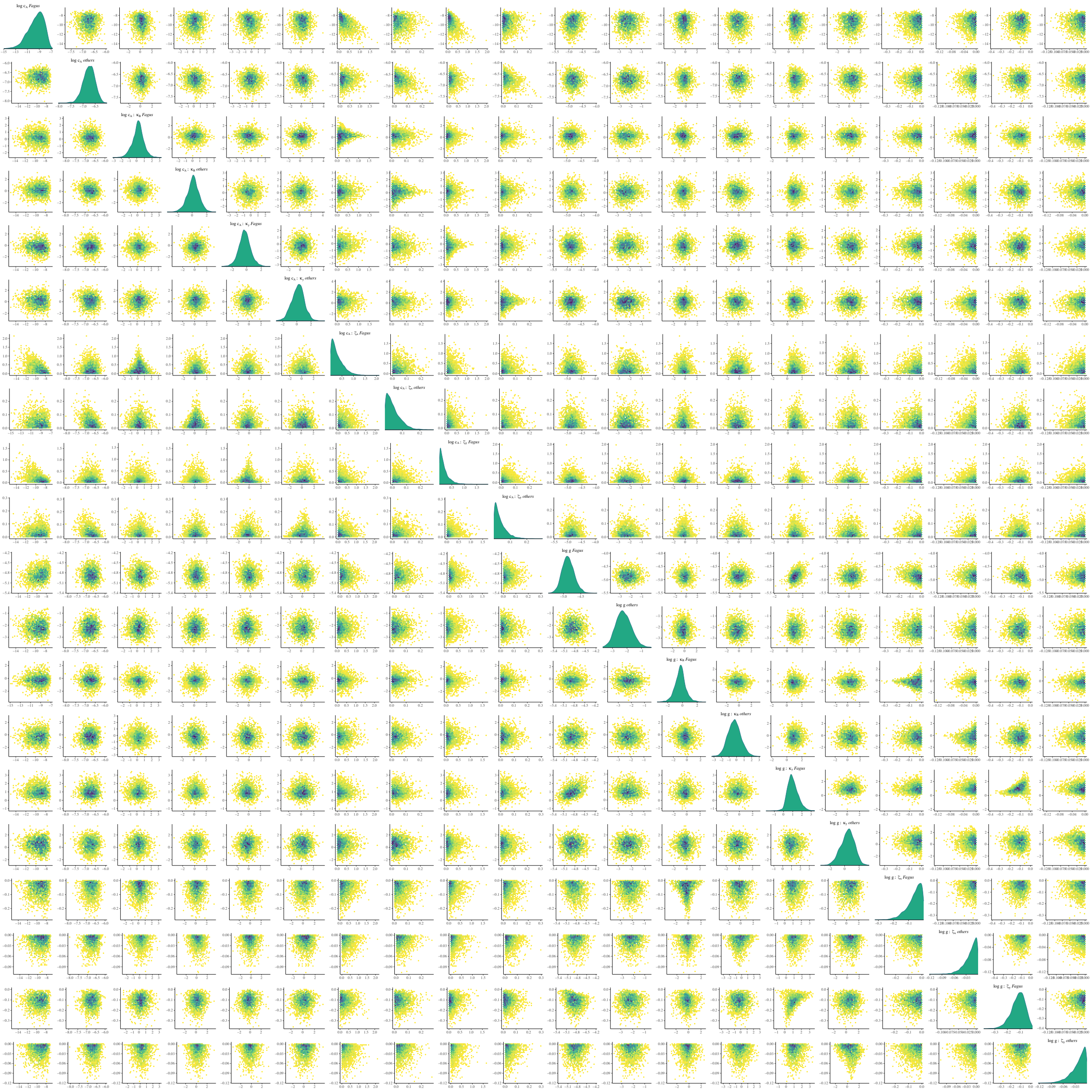

**Figure S5:** Pairs of paraboloid parameter correlations within stage A.

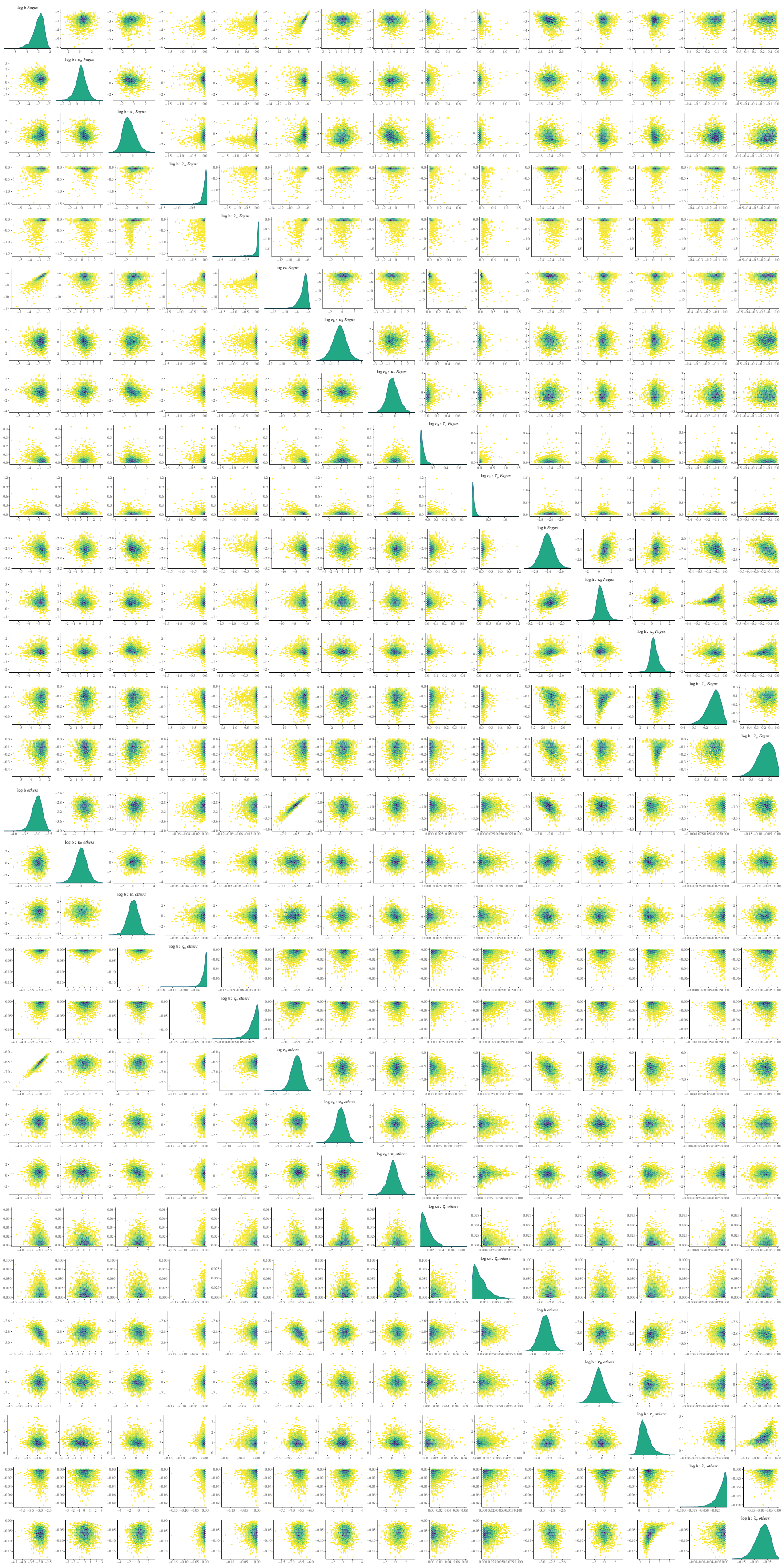

**Figure S6:** Pairs of paraboloid parameter correlations within stage B, separately for *Fagus* and *others*.

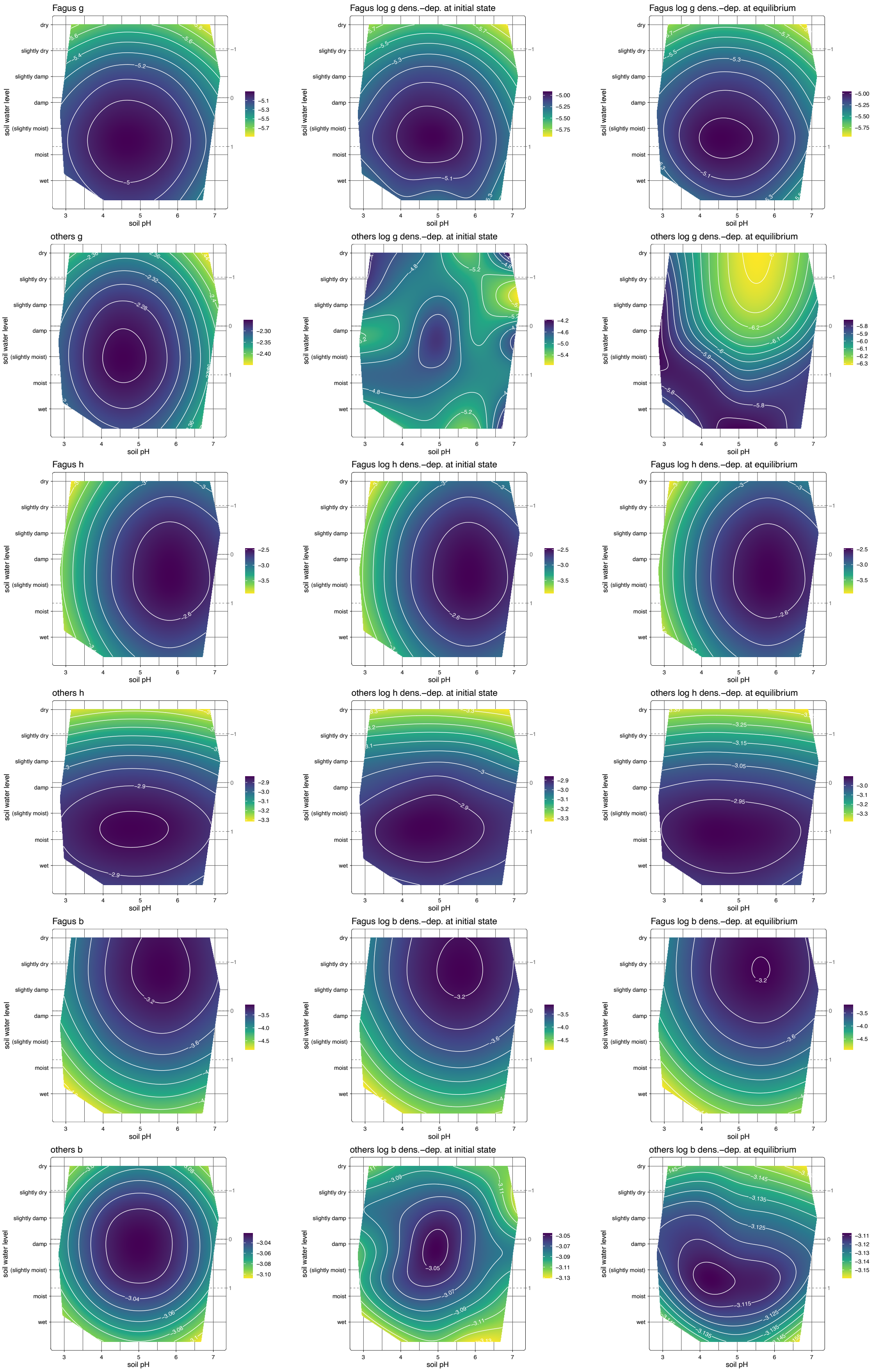


**Figure S7:** JAB model demographic rates in environmental space, including the density-dependent terms at initial and equilibrium state. The contours show predictions from smoothing splines fitted to the posterior distributions (as described in Section 2.5.1).

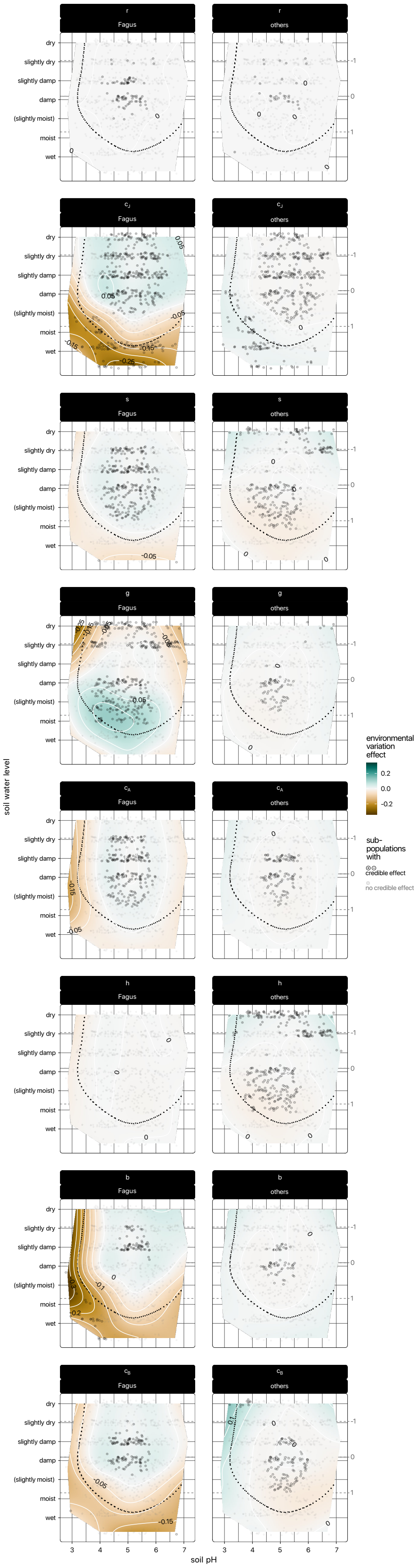


**Figure S8:** Effect of environmentally-responsive demography in environmental space for all rates that were determined with paraboloid parameters.

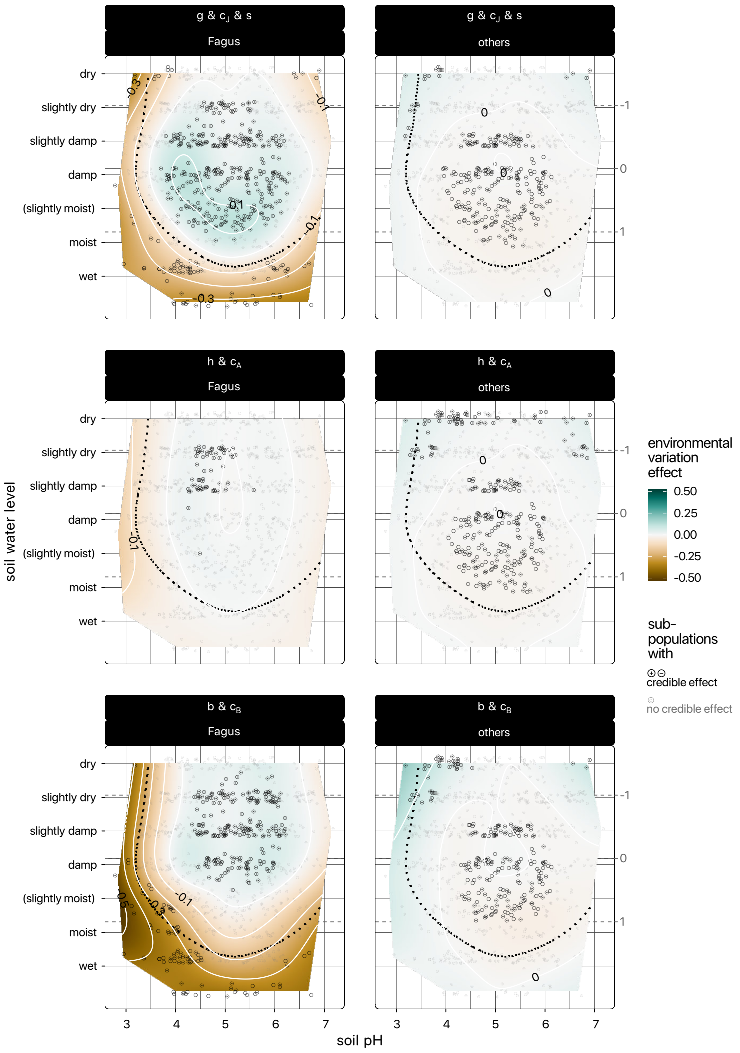


**Figure S9:** Effect of environmentally-responsive demography in environmental space for combinations of rates.

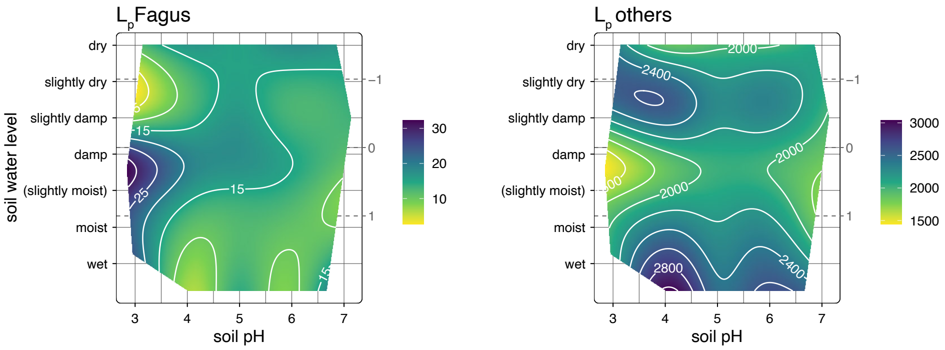


**Figure S10:** Effect of environmentally-responsive demography in environmental space for $L_p$.
